# *Legionella pneumophila* DenR is a bacterial S-acyltransferase for the human GTPase NRAS

**DOI:** 10.64898/2026.11.23.768205

**Authors:** Chad D. Williamson, Sabrina Bouchard, Ravi Pal, Stephanie S. Lehman, Gianlucca G. Nicastro, L. Aravind, Matthias P. Machner

## Abstract

In eukaryotic cells, the covalent attachment of palmitic acid to cysteine residues is catalyzed by the DHHC superfamily of membrane-embedded protein acyltransferases (DHHC-PAT). Here, we discovered that the bacterial pathogen *Legionella pneumophila* codes for its own protein acyltransferase, DenR, which hijacks host palmitoylation. Unlike the DHHC-PATs, DenR is a cytosolic protein that peripherally associates with membranes via binding of PI3P lipids through its carboxy-terminal domain. PI3P binding allosterically activates the palmitoyltransferase activity of its N-terminal circularly permuted NlpC/P60 papain-like domain, resulting in palmitoylation of the host GTPase NRAS at cysteine 181 and thereby anchoring NRAS to membranes of the *Legionella*-containing vacuole. Importantly, DenR-mediated S-acylation was insensitive to 2-bromopalmitate treatment, which inhibits cellular protein palmitoylation, suggesting that DenR represents a novel class of cytosolic S-acyltransferase.

**Significance Statement:** During infection, microbial pathogens often exploit cellular processes by encoding molecular mimics of host enzymes. Palmitoyltransferases are enzymes that catalyze palmitoylation, a form of protein lipidation that is conserved across all eukaryotes but has generally been considered absent in prokaryotes. Here, we show that the bacterium *Legionella pneumophila* utilizes palmitoylation during infection by encoding its own version of a palmitoyl transferase, the effector protein DenR. Unlike eukaryotic palmitoyltransferases, DenR is not an integral membrane protein but, instead, functions in the host cell cytosol where it catalyzes palmitoylation of NRAS, a small GTPase known for its role in cell growth and proliferation but also cancer. These findings suggest that palmitoyltransferases may have evolved independently at least twice during evolution.

## Introduction

All eukaryotic organisms encode members of the Ras-like family of small GTPases, sometimes tens to hundreds. Upon stimulation by upstream signals, Ras-like GTPases alternate between a GTP-bound active state and a GDP-bound inactive form, enabling them to function as molecular switches. Some of the most prominent members of the Ras superfamily in humans are NRAS and its three close paralogs, HRAS, KRAS4A and KRAS4B which regulate a diverse array of biological processes such as cell proliferation, differentiation and survival (1). Not surprisingly, gain-of-function mutations that result in constitutive Ras signaling contribute to nearly 30% of all human cancers, making these GTPases proto-oncoproteins.

Human Ras homologs have a nearly identical GTPase domain (G-domain) as well as a CAAX motif at their C-terminal end in which the cysteine (C) residue is the site of farnesylation, an irreversible form of lipidation that enhances membrane binding (2) (**Figure 1A**). Some RAS isoforms, such as NRAS and HRAS, receive a second type of lipidation called S-acylation, or palmitoylation, on either one (NRAS) or two (HRAS) cysteine residues within their C-terminal hypervariable region (HVR) upstream of the CAAX motif (**Figure 1A**). Palmitoylation further facilitates the interaction of NRAS or HRAS with membranes, allowing them to traffic via anterograde transport vesicles from the Golgi to the plasma membrane, where they eventually perform their signaling function (**Supporting Fig. 1A**). Since palmitoylation is a reversible form of lipidation, NRAS and HRAS undergo cycles of depalmitoylation and re-palmitoylation, which control their retrograde and anterograde transport, respectively, between the plasma membrane and the Golgi apparatus (**Supporting Fig. 1A)** (3, 4).

**Figure 1:**
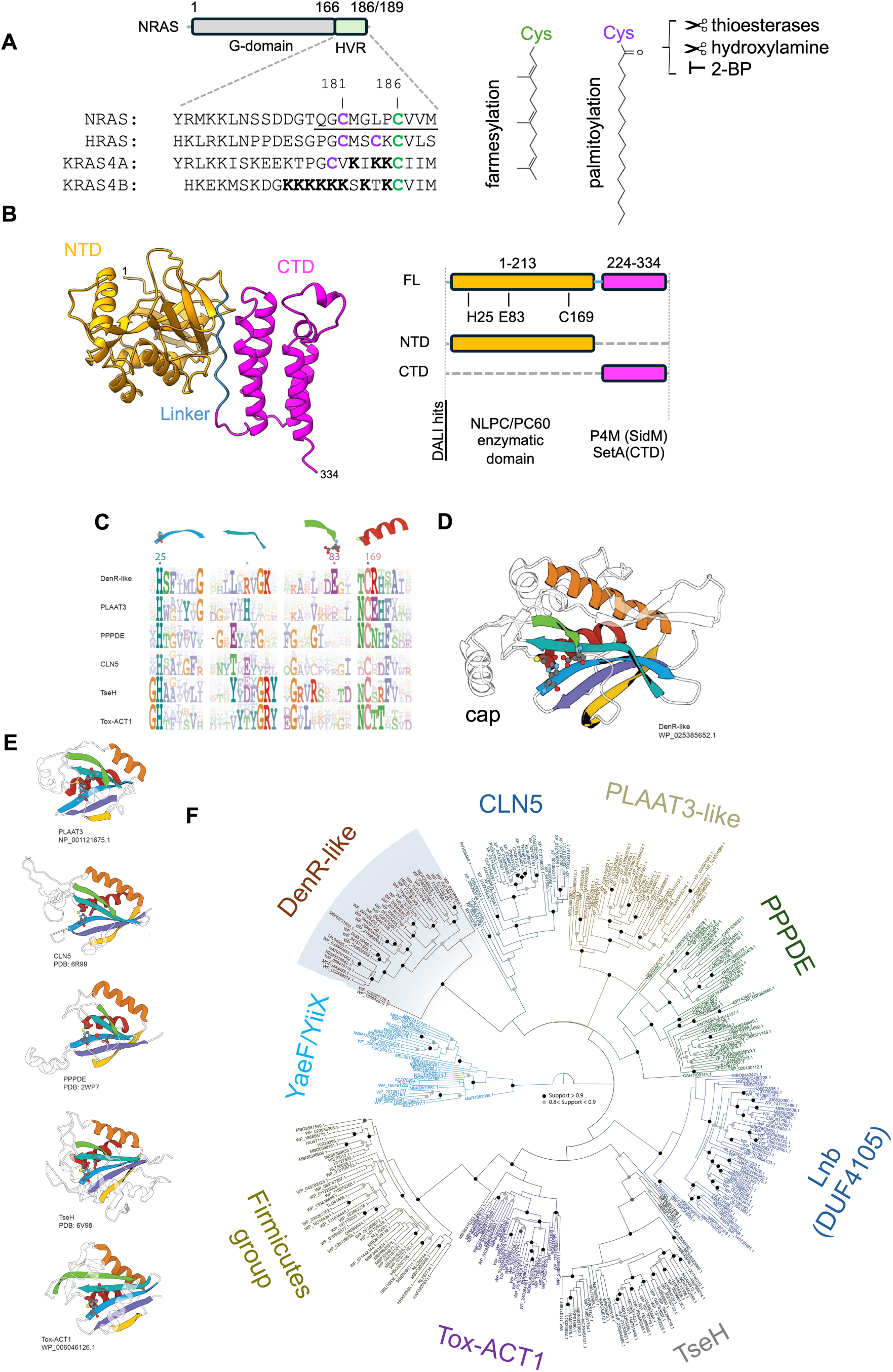
Diversity and evolutionary relationships of the permuted NlpC/P60 families. **(A)** The sequence of the hypervariable region (HVR) of NRAS is shown in comparison to that of the other RAS isoforms. Numbers indicate amino acid residues. Palmitoylated residues are in purple, farnesylated residues in green, positively charged residues in bold. Underlined residues are the sufficient for recruitment by DenR. Chemical structures of farnesylated vs palmitoylated cysteines and treatments that affect palmitoylation are shown on the right. **(B)** Ribbon diagram of DenR predicted by Alphafold 3. The N- and C-terminal domain are shown in yellow (NTD) and magenta (CTD), respectively. The poly-proline linker is colored in blue. Domain borders of truncated DenR variants and their closest structural homologs according to DALI are shown. **(C)** Sequence logo of the catalytic-site regions. The schematic above indicates the segment corresponding to the structures shown in (C, D). Black asterisks mark the nucleophilic cysteine and catalytic histidine. Blue asterisks denote the position of the third conserved residue (orienting residue) in the major families, whereas red indicates the alternative third residue characteristic of the DenR-like family. **(D)** Alphafold 3 model of the DenR-like NTD, with secondary structure elements color-coded and catalytic residues shown in ball-and-stick representation. **(E)** Structures of representative permuted NlpC/P60 family members. Catalytic residues are shown in ball-and-stick format. The red sphere in PPPDE denotes a water molecule coordinating the catalytic site. Experimentally determined structures are indicated by PDB identifiers; predicted models correspond to Alphafold 3 entries. **(F)** Phylogenetic tree depicting the evolutionary relationships among the distinct groups. Branches with support values >0.8 are marked by gray circles, and support values >0.9 are marked by black circles.

In eukaryotic cells, S-palmitoylation is catalyzed by a family of 23 protein acyltransferases (PATs) or DHHC enzymes, named after their conserved Asp-His-His-Cys catalytic motif (5–7). DHHC proteins localize to various organelle membranes where they catalyze the covalent attachment of a palmitoyl group (C16:0 fatty acid) or other long-chain acyl groups to the side chain of cysteine residues via a thioester bond. The thioester-linked acyl groups can be cleaved either enzymatically by thioesterases or chemically via hydroxylamine treatment (**Figure 1A**). While individual DHHC enzymes may differ in their subcellular localization, tissue expression pattern, molecular size, and domain organization, they all contain four to six transmembrane helices and are membrane-embedded, with their cysteine-rich region comprised of three partial treble-clef-like repeats that house the DHHC catalytic site facing the cytosol (8). Notably, DHHC enzymes are absent from prokaryotes, and bacteria capable of catalyzing protein palmitoylation have yet to be discovered.

In this report, we provide evidence for a bacterial pathogen that can utilize palmitoylation during infection by encoding its own version of a palmitoyltransferase.

## Results

### The DenR N-terminal domain (NTD) represents a novel version of the circularly permuted NlpC/P60 clade

*Legionella pneumophila* is the causative agent of Legionnaires’ pneumonia in humans. During infection of human cells, the *Legionella*-containing vacuole (LCV) becomes decorated with NRAS (9). NRAS colocalization with LCVs is detectable as early as 30 minutes after bacterial uptake and progressively intensifies until most NRAS is depleted from the host cell plasma membrane, its normal site of signaling (9). As a result, signaling along several host pathways, including the MAP kinase pathway, was either modulated or dampened (9). We also showed that NRAS recruitment to the LCV depends on DenR (Lpg1909), an effector protein that is delivered into host cells via the *Legionella* Dot/Icm type IV secretion system (T4SS) (9). Notably, DenR-mediated recruitment was specific for NRAS, while its homologs HRAS and KRAS4A/4B were absent from LCVs (9).

To explore the molecular mechanism of NRAS recruitment by *L. pneumophila*, we performed *in silico* structural analyses with DenR. The highest-scoring Alphafold 3 model predicted two distinct domains for DenR from *L. pneumophila* strain Philadelphia-1, hereafter termed N-terminal domain (NTD) and C-terminal domain (CTD), connected by a flexible linker rich in proline residues (**Figure 1B; Supporting Fig. 1B**). DenR is encoded by more than 80% of *Legionella* species (36 out of 41 genomes (10); 65 out of 80 genomes (11)) and the closely related pathogen Fluoribacter. Despite considerable sequence variability among the homologs (98% to 26% identity) (**Supporting Fig. 1C**), their predicted structures were remarkably similar (**Supporting Fig. 1D)**. A profile-profile search using the DenR(NTD) against a custom library of profiles of domains found in bacterial effectors and other toxins yielded hits to several members of the vast NlpC/P60 clade of the papain-like superfamily. This was confirmed by DALI searches against the PDB database (**Figure 1B**, right) that yielded hits with high Z-scores (>6) to several representatives of the NlpC/P60 clade, including acyltransferase/hydrolase enzymes that act as peptidases, acyl hydrolases (including lipases) and transacylases acting on amide and ester linkages.

The conserved core of the Papain-like fold in the NlpC/P60 clade is characterized by two α-helices and 5 strands forming a β-sheet, with a “V”-shaped split between strands 2 and 3. The active site is a characteristic catalytic triad (12), housed in the split in the β-sheet and the helix appressed against it (**Figure 1C-E**). In the predicted DenR(NTD) structure, the split β-sheet bears the catalytic histidine (H25) and a second polar residue (E83), whereas the beginning of the appressed helix bears the catalytic cysteine (C169) (**Figure 1C**). Notably, in the ancestral version of the papain-like fold, the helix bearing the invariant catalytic cysteine is located at the N-terminus of the domain **(Supporting Figure 1E)**; however, in DenR and other circularly permuted versions of the NlpC/P60 clade the permutation moves it to the C-terminus of the β-sheet while retaining an intact overall 3D structure (**Figure 1C, D; Supporting Figure 1E**). The third residue of the catalytic triad in the DenR branch is a glutamate (E83) just downstream of strand 4 **(Figure 1C)**. Another notable feature in several branches of the NlpC/P60 clade is an insert between strand 3 and 4 of the core sheet which forms a “cap” over the active site (**Figure 1D, E**). DenR is predicted to also possess an equivalent cap which is more rapidly evolving than the rest of the catalytic domain suggesting a role in either guarding the active site from host immunity factors or facilitating substrate recognition/binding.

A phylogenetic analysis was performed that included several newly identified branches of the circularly permuted NlpC/P60 clade that have not previously been included in such a study. The resulting tree revealed that the DenR branch is a distinct lineage that is deeply lodged within the circularly permuted clade with the eukaryotic depalmitoylating enzyme CLN5 as its immediate sister group, followed by the PLAAT-like and PPPDE branches (**Figure 1F**). Both of these and more distant branches of the circularly permuted clade, such as TseH, the ACT toxin, and Lbn (13) have been shown or predicted to act on lipids and tend to have a hydrophobic environment surrounding the active site residues, which is also preserved in the DenR branch. These findings suggested a potential role for the DenR(NTD) in catalytically acting on lipids or long-chain fatty acyl linkages.

### A catalytic triad within the N-terminal domain is required for NRAS recruitment by DenR

Our earlier findings had demonstrated that the C-terminal ten amino acids of the HVR of NRAS (underlined in **Figure 1A**), including C181 and C186 which are the targets of palmitoylation and farnesylation, respectively, were entirely sufficient for recruitment by DenR (9). When Alphafold 3 was tasked to assemble a heterotrimeric complex of DenR, C186-lipidated NRAS, and free palmitic acid, its highest confidence model positioned the HVR of NRAS into the putative active site of DenR, with C181 on top of the catalytic triad while free palmitic acid reached into the characteristic hydrophobic pocket of the circularly permuted NlpC/P60 from the other side so that both substrates come close enough for S-acylation of C181 to occur (**Figure 2A**). Upon substituting the putative catalytic residues with alanine, we found that none of the DenR variants showed substantial colocalization with GFP-NRAS (**Figure 2B**), while wild-type mCherry-DenR showed the previously described robust accumulation on large vesicular structures where it colocalized with GFP-NRAS (**Figure 2B**) (9). Quantitative analyses indicated that, of the catalytic site mutations, H25A and C169A had the most severe effect on NRAS recruitment (**Figure 2B**, graph). Notably, substitution of an unrelated NTD residue (L68, situated in the predicted cap region) with alanine did not abrogate DenR’s ability to recruit NRAS (**Supporting Fig. 2A)**.

**Figure 2:**
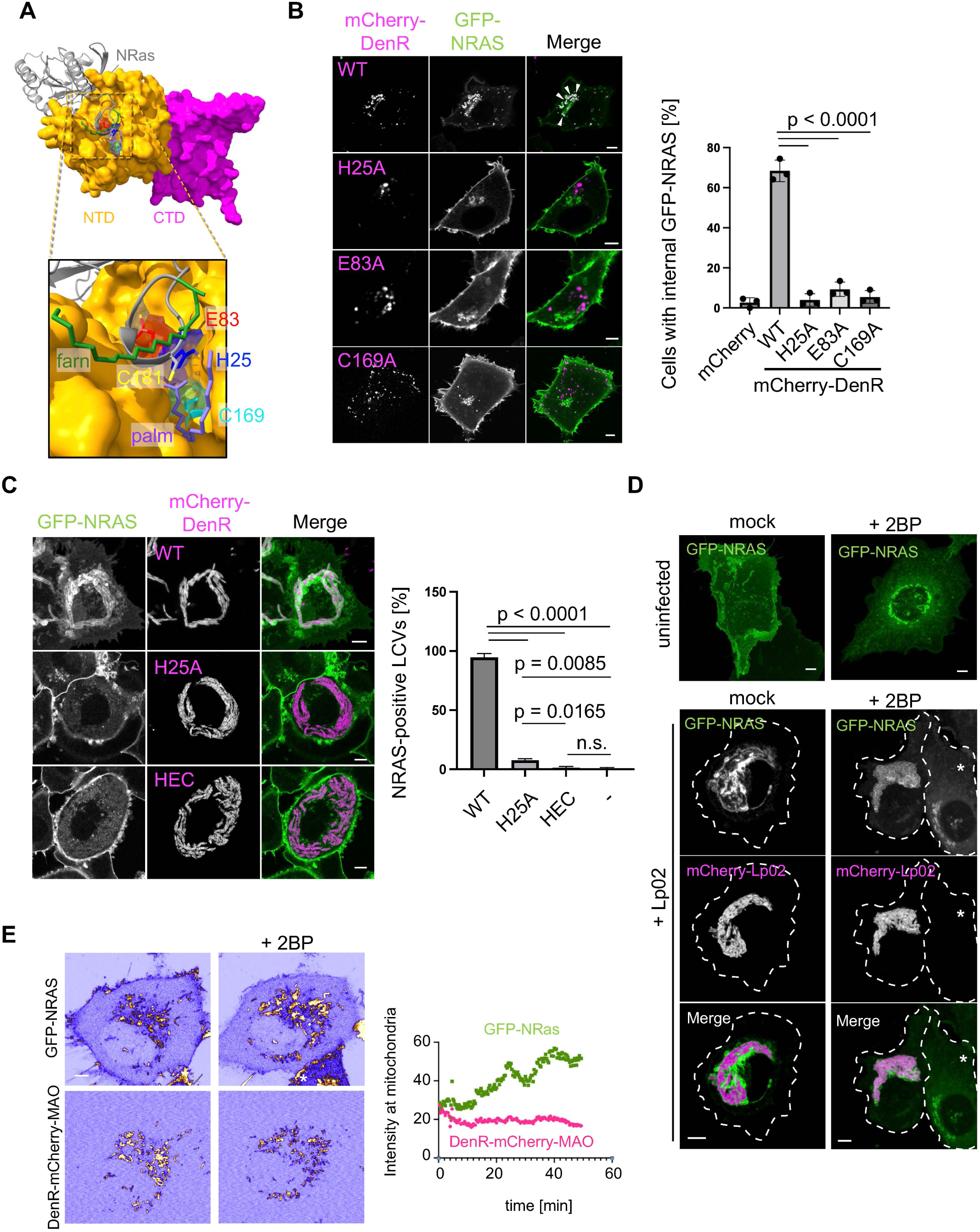
A catalytic triad within the NTD is required for NRAS recruitment, independent of host palmitoyl transferases. **(A)** Alphafold 3 model of DenR shown as surface representation in complex with palmitic acid and Rce1-cleaved NRAS (ribbon model) lipidated at C186. The inset shows a magnified view of the active site. The catalytic residues H25, E83, and C169 are shown as sticks and are color-coded. The highest confidence model positioned the HVR of NRAS (grey) into the catalytic pocket, with C181 and free palmitic acid (purple) in proximity to the catalytic residues for S-acylation of C181 to occur. Note that the lipidated C186 (we used palmitoylation since Alphafold 3 did not offer a farnesylation option) at the C-terminal end of the HVR is positioned in a way that allows for it to potentially rest in a membrane while C181 is modified with palmitic acid. **(B)** Active site mutants of DenR fail to recruit NRAS to endomembranes. Transiently transfected HT1080 cells producing GFP-NRAS and the indicated mCherry-DenR variants were imaged by fluorescence microscopy. Arrowheads indicate examples of areas with NRAS-DenR colocalization. The graph represents quantification of three experiments (n=3). **(C)** Active site mutants of DenR fail to recruit NRAS to LCVs. HT1080 cells producing GFP-NRAS were challenged with Lp02Δ*denR* complemented with plasmids encoding the indicated DenR variants, and GFP-NRAS colocalization with LCV was examined 16 hpi by fluorescence microscopy. Graph represents quantification of three experiments (n=3). **(D)** NRAS localization to LCVs is unaffected by 2BP. HT1080 cells producing GFP-NRAS were challenged with mCherry-producing Lp02 (bottom panels), or left uninfected (top panels), and imaged with or without overnight incubation with the DHHC enzyme inhibitor 2BP (25 mM). Images are maximum intensity projection micrographs. The cell periphery of infected cells is indicated by dashed lines; asterisks indicate uninfected bystander cells where 2BP treatment causes NRAS localization to the Golgi. **(E)** Inhibition of cellular DHHC enzymes stimulates NRAS recruitment by DenR. Transiently transfected HT1080 cells producing DenR-mCherry-MAO and GFP-NRAS were treated with 2BP (75 mM), and recruitment of NRAS to DenR-positive mitochondria was examined by live fluorescence microscopy for 49 min (micrographs from the start and end of the imaging series are displayed, pseudocolored using the ICA lookup table to better highlight changes in intensity). The graph quantifies NRAS and DenR fluorescence intensity at mitochondria over time upon 2BP treatment in the representative cell displayed. All scale bars, 5 µm.

Catalytic site mutants of DenR still interacted with NRAS in co-precipitation assays, indicating that the mutations did not cause general defects in protein folding or stability (**Supporting Figure 2B**). DenR(H25A) or a DenR triple mutant (H25A, E83A, C169A; DenR(HEC)) also failed to rescue the NRAS recruitment defect of a Δ*denR* strain 6 hours post infection (hpi) (**Supporting Fig. 2C**) or 16 hpi (**Figure 2C**). Together, these data are consistent with the idea that DenR(NTD) recruits host cell NRAS to LCVs through its catalytic activity, possibly via C169-dependent palmitoylation.

### NRAS recruitment by DenR is independent of host palmitoyltransferases

Our previous work had demonstrated that replacement of cysteine at position 181 in NRAS with serine prevented its recruitment to LCVs (9). C181 is known to be the target of reversible palmitoylation by cellular DHHC proteins (**Figure 1A**). We assayed how NRAS recruitment to LCVs is altered in the presence of 2-bromopalmitate (2BP), an irreversible inhibitor of palmitoylation by cellular DHHC enzymes (14). As expected, 2BP treatment of uninfected HT1080 cells efficiently blocked palmitoylation of NRAS by cellular DHHC proteins, resulting in GFP-NRAS relocating from the plasma membrane to the Golgi compartment (**Figure 2D**; uninfected). When HT1080 cells producing GFP-NRAS were challenged with mCherry-producing Lp02, NRAS showed robust colocalization with LCVs even under conditions of 2BP treatment (**Figure 2D**; +Lp02), while uninfected bystander cells showed the previously reported relocation of NRAS from the plasma membrane to the Golgi (**Figure 2D**, star). We also tested the effect of 2BP treatment on NRAS recruitment to mitochondria in HT1080 cells producing DenR-mCherry conjugated to a mitochondria-targeting sequence (MAO), which directs proteins to the cytoplasmic face of the outer mitochondrial membrane (OMM), a location where NRAS is not typically found. Similar to the results from the infection experiment (**Figure 2D**), acute 2BP treatment did not inhibit NRAS recruitment to mitochondrial membranes by DenR-mCherry-MAO (**Figure 2E**); if anything, it increased it by nearly two-fold. No mitochondrial enrichment of NRAS was observed in HT1080 cells lacking DenR-mCherry-MAO and treated with 2BP (**Supporting Fig. 2E**). These data suggest that the catalytic activity of DenR was insensitive to 2BP and that NRAS recruitment by DenR was not dependent on cellular DHHC enzymes.

### DenR is a bacterial palmitoyltransferase

To determine the palmitoylation state of NRAS in the presence of DenR, we utilized an acyl resin-assisted capture (Acyl-RAC) assay that, under conditions of H_2_NOH treatment, enriches palmitoylated proteins. In the absence of DenR, only background levels of NRAS binding to thiopropyl sepharose beads were observed (**Figure 3A**). However, in the presence of DenR, a dramatic increase in NRAS capture by thiopropyl sepharose beads was detected with addition of H_2_NOH (**Figure 3A**), indicating that the DenR-catalyzed reaction on NRAS was indeed S-acylation.

**Figure 3:**
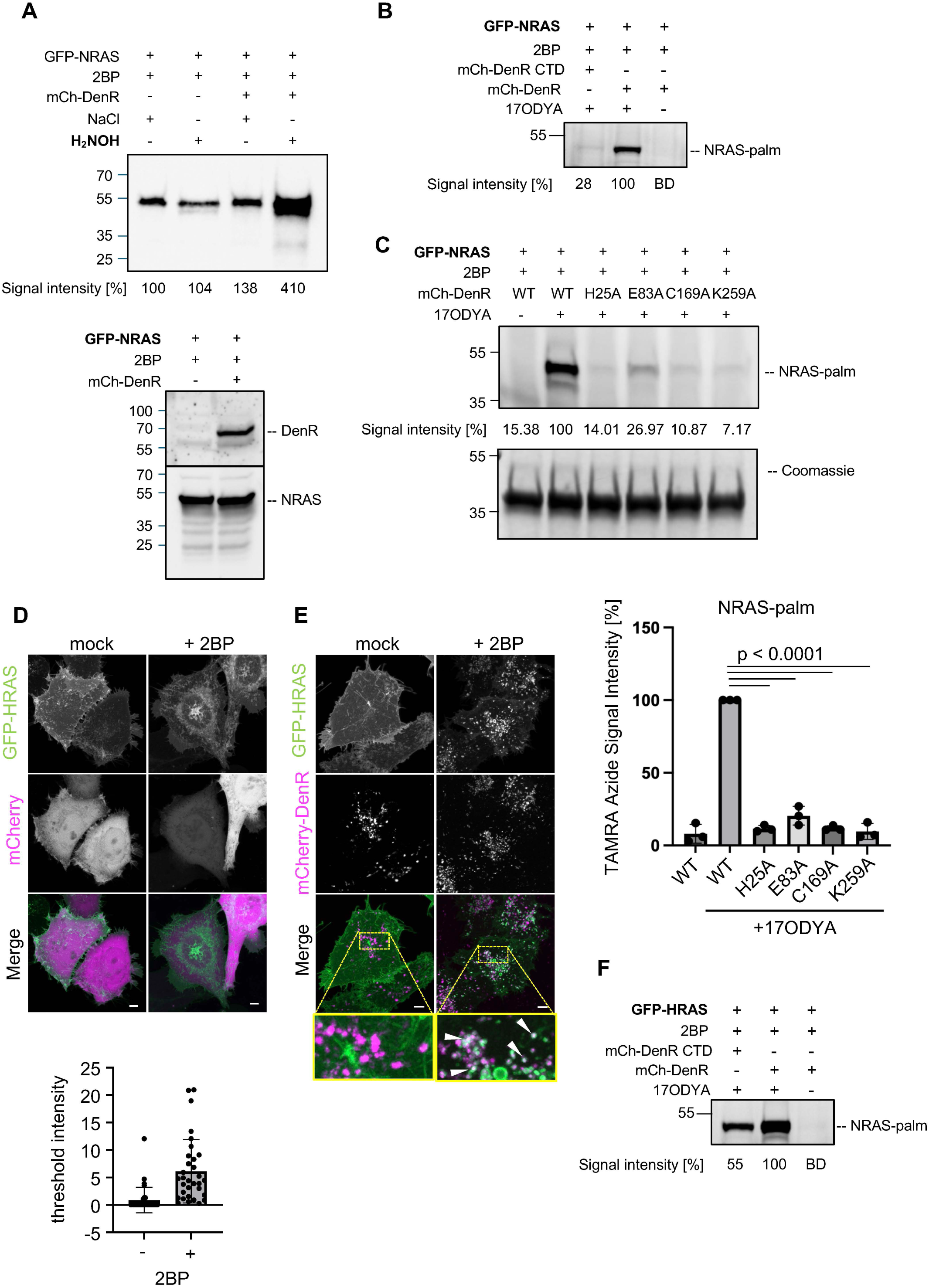
DenR has S-acyltransferase activity. (**A**) Acyl-RAC assay to identify thioester bond formation within NRAS. Transiently transfected HEK293T cells producing GFP-NRAS alone, or co-producing GFP-NRAS and mCherry-DenR, were treated with 2BP (25 mM) overnight to inhibit palmitoylation by DHHC enzymes. Thiopropyl sepharose beads were then used to capture S-acylated proteins from cell lysates treated either with hydroxylamine (H2NOH) or NaCl, a control condition to reveal background binding level of proteins with beads. Numbers indicate band intensities relative to control (lane 1; arbitrarily set to 100%). Bottom panel shows expression levels of GFP-NRAS and mCherry-DenR in cell lysates. (**B, C**) Click chemistry to detect the lipidation of proteins. 2BP-treated HEK293 cells producing the indicated DenR variants and GFP-NRAS were metabolically labelled with 17ODYA. GFP-tagged NRAS was affinity-purified by immunoprecipitation with GFP nanobody beads, and palmitoylation was detected by click chemistry with TAMRA-azide and in-gel visualization using a ChemiDoc Imaging System. Numbers indicate signal intensity relative to WT DenR with 17ODYA which was arbitrarily set to 100%. BD, below detection. Similar click chemistry experiments were carried out as above to compare activity of DenR(WT) to indicated DenR point mutants on NRAS acylation (C). Graph below represents quantification of three experiments (n=3), mean +/− SD. (**D, E**) Inhibition of cellular DHHC enzymes makes HRAS accessible for recruitment by DenR. Transiently transfected HeLa cells producing GFP-HRAS and either mCherry (D) or mCherry-DenR (E) were treated with 2BP, and recruitment of HRAS to endomembranes was examined by fluorescence microscopy. Insets represent examples of endomembranes; white arrowheads indicate regions of colocalization. The graph below is an intensity quantification of GFP-HRAS signal recruited to DenR-positive membranes for at least 30 cells per condition from a single representative experiment (E). (F) Same experiment as (B) but with GFP-HRAS. All scale bars, 5 µm.

17-octadecynoic acid (17-ODYA) is a palmitic acid analog used in copper-catalyzed azide-alkyne cycloaddition reactions (click chemistry) to detect the lipidation of proteins. We metabolically labeled transiently transfected HEK293 cells with 17-ODYA and used click chemistry to detect DenR-mediated lipidation of NRAS in the presence of 2BP. Robust lipidation of GFP-NRAS was observed only in cells producing full-length DenR (mCherry-DenR) but not in cells producing mCherry-DenR(CTD), which lacks the NTD, or in cells lacking 17-ODYA (**Figure 3B**).

Next, we examined the effect of catalytic site mutations on the ability of DenR to modify GFP-NRAS (**Figure 3C**) with 17-ODYA. Click chemistry assays using lysate from HEK cells revealed that DenR(H25A) and DenR(C169A) were the catalytic site variants most strongly attenuated in lipidating NRAS (∼88% reduction in signal compared to WT). These data matched the results of the cell-based recruitment assays, where mCherry-DenR(H25A) and mCherry-DenR(C169A) showed the lowest level of colocalization with GFP-NRAS (**Figure 2B**). We similarly utilized click chemistry assays to examine the ability of DenR to form reaction intermediates with 17-ODYA, a known mechanistic feature of DHHC enzymes. 17-ODYA labeling was readily observed for mCherry-DenR(WT) within HEK cell lysate, while catalytic site mutations significantly inhibited DenR acylation of the catalytic cysteine (**Supporting Fig. 3A**). Together, these data confirm that DenR is an S-acyltransferase that forms a transient acyl-enzyme intermediate prior to transferring the lipid moiety to free cysteines within the HVR of NRAS.

### DenR preferentially recruits de-palmitoylated Ras GTPases

Unlike NRAS, wild-type HRAS was not recruited to LCVs (9) despite the fact that it contains cysteine residues within its HVR, Cys181 and Cys184, that are the sites of palmitoylation by cellular DHHC enzymes (**Figure 1A)**. On the other hand, HRAS(C181S), which lacked one of its palmitoylation sites, was abundantly present on LCVs (9). HRAS(C181S) is known to have a more pronounced presence at the Golgi than HRAS WT or C184S (15), suggesting that Golgi localization may be a determining factor for recruitment by DenR. To further test this hypothesis, we produced GFP-HRAS in HT1080 cells with either mCherry or mCherry-DenR, followed by treatment of cells with 2BP to inhibit cellular DHHC enzymes, thus shifting the equilibrium towards the depalmitoylated state of HRAS. In control cells producing mCherry, GFP-HRAS was abundantly present at the plasma membrane and, upon 2BP treatment, retreated to the Golgi (**Figure 3D**). In cells producing mCherry-DenR (**Figure 3E**), GFP-HRAS also disappeared from the plasma membrane upon 2BP treatment but, strikingly, the protein was now enriched on internal, DenR-positive endomembranes. Similar results were obtained with GFP-HRAS(C184S) (**Supporting Fig. 3B, C**). Notably, not all proteins with a palmitoylation site were a target of DenR. The *L. pneumophila* effector GobX localized to Golgi membranes via host-mediated palmitoylation at a cysteine residue within its C-terminal region (amino acids 171-190; GFP-GobX_171-190_) (**Supporting Fig. 3D,** top row) (16) but lost this Golgi enrichment upon incubation with 2BP (second row). In the presence of mCherry-DenR, GFP-GobX_171-190_ did not relocate to DenR-positive membranes regardless of 2BP treatment (**Supporting Fig. 3D**, bottom rows), consistent with the idea that recruitment by DenR is a selective process where prolonged residency at the Golgi is beneficial but not sufficient.

Lastly, we performed click chemistry assays with HRAS under conditions of 2BP treatment. Similar to NRAS, HRAS was robustly modified with 17-ODYA in lysate from transiently transfected HEK cells producing DenR WT upon 2BP treatment, while 17-ODYA incorporation was dramatically reduced in cells producing DenR(CTD) (**Figure 3F**) or cells producing catalytic mutants DenR(H25A), DenR(E83A) or DenR(C169A) (**Supporting Fig. 3E**).

### The CTD of DenR is sufficient for membrane binding but not NRAS recruitment

A profile-profile search with DenR(CTD) using HHPred (17) revealed high-scoring hits (probability 80-90%) to the profiles of the P4M domain found in SidM (DrrA; Lpg2464, PDB 4MXP; (18)) and SetA (Lpg1978; 8X4N; (19)) (**Figure 4A)**. These structures were also recovered as the best-scoring hits in a DALI search against the PDB database using the Alphafold 3-predicted structure of the DenR(CTD) (**Figure 1B**). The DenR(CTD) shared a 4-helical core with the P4M domain, along with an extended loop with a short helical segment between the core helices 3 and 4, which forms a phospholipid-headgroup-binding site. Further, this four-helical core of the P4M domain also resembles the equivalent core of the membrane localization domain (MLD) found in several effectors/toxins from *Legionella* species and other bacteria (e.g., *Pasteurella multocida*, PDB: 2EBH, (20)) (**Supporting Fig. 4A,B**) (21–24). Together, these observations suggested that the DenR(CTD), like other P4M domains, might localize the protein to target membranes. We generated plasmids encoding either full-length (FL) DenR or truncated DenR variants (NTD or CTD) conjugated to green fluorescent protein (GFP). Upon production in transiently transfected HeLa cells (**Figure 4B**), the P4M-like DenR(CTD) displayed a localization pattern similar to that of GFP-DenR(FL), suggesting that the CTD had retained membrane binding capability. In contrast, DenR(NTD) showed a diffuse cytosolic fluorescence signal and was not enriched on membranes (**Figure 4B**).

**Figure 4:**
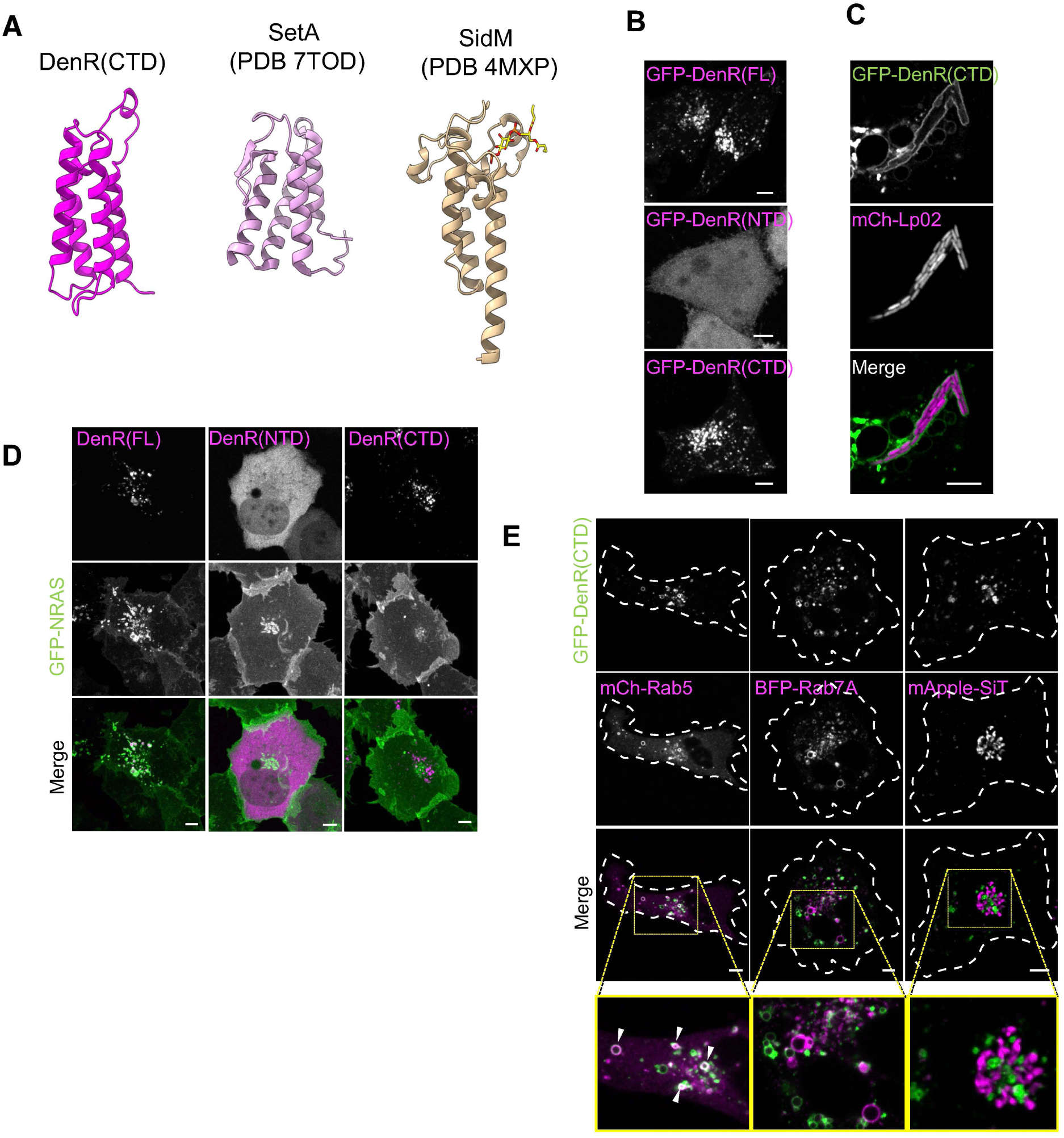
DenR CTD has membrane localization capabilit. **(A)** Ribbon diagram of Alphafold 3-predicted structure of DenR(CTD) compared to experimentally determined structures of its closest structural homologs (PDB database identifiers are indicated). The lipid binding site in SidM-P4M is bound to the PI4P analog DB4 shown as stick model. **(B)** DenR(CTD) is required for localization to endomembranes. Images show representative maximum intensity projection examples of HeLa cells producing the indicated DenR variants. **(C)** The CTD is sufficient for LCV localization. HT1080 cells producing GFP-DenR(CTD) were challenged with mCherry-producing *L. pneumophila* (Lp02) for 12 hours, and localization of GFP-DenR(CTD) to LCVs was examined by live microscopy. Images show confocal slices of Airyscan super-resolution images of the LCV. **(D)** The CTD alone is not sufficient to recruit NRAS to membranes. GFP-NRAS and the indicated mCherry-DenR variants were co-produced in HT1080 cells, and their localization was examined by live microscopy. Images show maximum intensity projections of representative cells. **(E)** DenR(CTD) targets to endosomal compartments. GFP-DenR(CTD) was co-produced with the indicated organelle markers in HT1080 cells, and their localization was examined microscopically. Insets represent examples of endomembranes; white arrowheads indicate regions of colocalization. All scale bars, 5 µm.

To further test if the DenR(CTD) was able to recognize membranes of the LCV, transiently transfected HT1080 cells producing GFP-DenR(CTD) were challenged with *L. pneumophila* wild-type (Lp02) for 12 hours and then imaged by live microscopy (**Figure 4C**). GFP-DenR(CTD) was clearly detectable on the surface of the LCV, confirming that the CTD was sufficient for DenR’s membrane-targeting activity and its localization to LCVs during infection.

Next, we examined whether DenR(CTD) could alter the localization of NRAS in the absence of the NTD. We performed live microscopy on HT1080 cells producing both GFP-NRAS and either mCherry-DenR(FL) or mCherry-DenR(CTD) (**Figure 4D**). As reported before (9), in the absence of LCV membranes, DenR(FL) was localized to large vesicular organelles, and it recruited GFP-NRAS to their membranes (**Figure 4D**; left panels). In contrast, DenR(CTD), despite localizing to similar endomembranes, was unable to redirect GFP-NRAS from the plasma membrane to those organelles (right panels). As expected, mCherry-DenR(NTD) with its cytosolic distribution pattern did not alter NRAS localization (middle panels). These data suggest NRAS capture by DenR requires both membrane localization and catalytic activity.

To determine the organelle preference of recombinant DenR(CTD), we compared its localization to that of markers of early or late endosomes, namely Rab5 or Rab7, as well as the trans-Golgi network marker sialyltransferase (SiT) (**Figure 4E**). While little to no colocalization of GFP-DenR(CTD) was observed with blue fluorescent protein (BFP)-tagged Rab7 or mApple-SiT, there was substantial colocalization with the early endosomal marker mCherry-Rab5 in uninfected cells where LCVs are absent (**Figure 4E**).

### The C-terminal domain of DenR is a phospholipid binding module

Phosphoinositide (PI) lipids are important signaling molecules generated via phosphorylation of their inositol headgroup at the 3′, 4′, and 5′ position. The P4M domain of SidM has high selectivity for PI4P lipids which are typically found at Golgi membranes but also membranes of the early LCV (25). A critical lysine residue at position 568 in P4M (K568; highlighted in dark gray) engages the 4-phosphate group of PI4P (**Figure 5A)**, and its substitution with alanine strongly attenuated PI4P binding (18). Interestingly, even though the DenR(CTD) sequence was highly divergent from the P4M domain of SidM and belonged to a distinct clade found in numerous *Legionella* effectors (**Supporting Fig. 4B**), it contained a lysine residue (K259) at the equivalent position within the predicted phospholipid binding pocket (**Figure 5A**).

**Figure 5:**
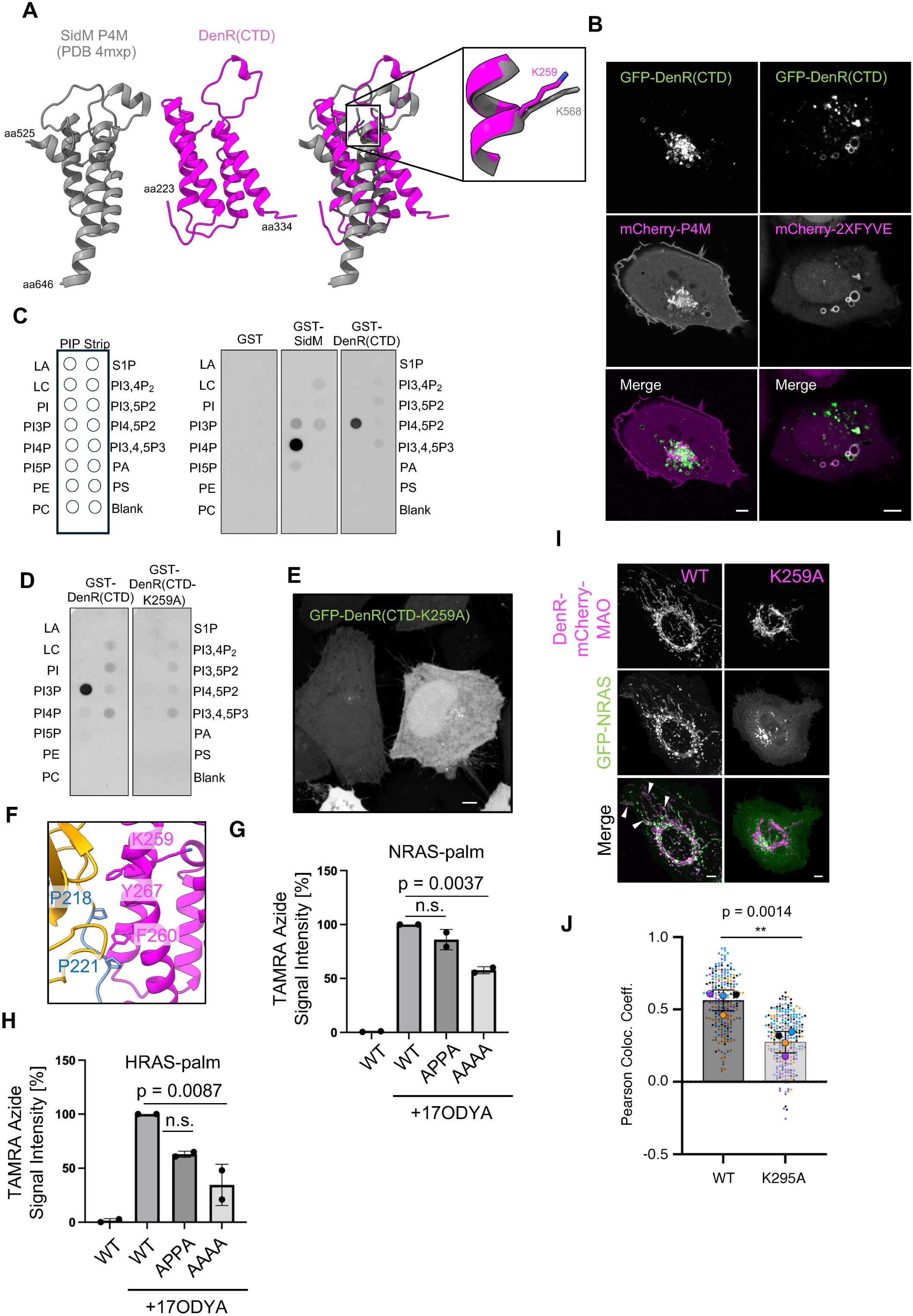
DenR CTD is a PI3P-specific membrane binding module. **(A)** Side by side comparison of the C-terminal domains of SidM (left, experimentally determined structure) and DenR (middle, predicted structure) shown as ribbon diagram, with a superposition shown on the right. The conserved lysine residue is represented as stick in the inset. **(B)** DenR(CTD) localizes to PI3P-positive membranes. GFP-DenR(CTD) was coproduced in HT1080 cells with either mCherry-P4M or mCherry-2xFYVE and examined microscopically 24 hours after transfection. **(C)** DenR(CTD) binds PI3P. Purified GST-SidM or GST-DenR(CTD) were applied to PIP strips, and lipid binding was determined using antibody-based detection. GST was used as a control. The template indicating the position of each type of lipid on the PIP strip is shown on the left. **(D)** The conserved lysine residue K259A is required for PI3P binding. Top: Same experiment as in (C) comparing GST-DenR(CTD) WT vs K259A. **(E)** Fluorescence maximum intensity projection micrograph showing failure of GFP-DenR(CTD-K259A) to localize to endomembranes in transiently transfected HeLa cells. **(F)** Ribbon diagram of DenR with conserved residues predicted to relay PI3P binding by the CTD to the NTD shown as sticks. (**G,H**) Click chemistry to detect the role of conserved linker residues on substrate lipidation. Click chemistry experiments were performed as in Figure 3C and Supporting Figure 3E to compare mutations in conserved linker residues (F) in lipidation of NRAS (G) or HRAS (H) substrates. Graphs show mean +/− SD quantification of TAMRA-azide signal detected from two experiments (n=2). Representative blots are shown in Supporting Figure 5C,D. **(I)** NRAS recruitment by DenR requires PI3P binding. Transiently transfected U2OS cells producing GFP-NRAS and either DenR-mCherry-MAO WT or the K259A mutant were imaged by fluorescence microscopy. Arrowheads indicate NRAS on mitochondria enriched in DenR WT but not the PI3P binding mutant. **(J)** NRAS recruitment by DenR requires PI3P binding. Same experiment as (G), but in HEK cells (micrographs shown in Supporting Fig. 5E). Graph shows quantification of pearson colocalization coefficient between mCherry and GFP signals per cell plotted as a superplot. Independent experiments (n=4) are color-coded with small dots representing individual cells and large dots representing the average for each experiment. ** = (p < 0.01) All scale bars, 5 µm.

Unexpectedly, co-production of GFP-DenR(CTD) and mCherry-P4M in transiently transfected HT1080 cells revealed little to no colocalization between the two proteins (**Figure 5B)**, suggesting that they localize to different cell compartments. We did, however, see substantial colocalization of DenR(CTD) with mCherry-2XFYVE, a probe for PI3P lipids, which are enriched on endosomes (**Figure 5B**).

To determine the exact phospholipid preference of DenR(CTD), we performed lipid-protein overlay assays. Purified glutathione S-transferase (GST)-tagged SidM or GST-DenR(CTD) were applied to lipid-coated nitrocellulose membranes (PIP strips), followed by antibody-mediated detection (**Figure 5C**). While GST-SidM exhibited the previously reported preference for PI4P (25), GST-DenR(CTD) showed robust binding to PI3P (**Figure 5C**) which was abolished upon substituting the conserved K259 with alanine (**Figure 5D**). GST (control) showed no notable binding to any of the immobilized lipids. Likewise, GFP-DenR(CTD-K259A) was unable to localize to endosomal membranes when produced within transiently transfected HeLa cells (**Figure 5E**). Together, these data establish DenR(CTD) as a P4M-like domain with membrane targeting activity and confirm the critical role of K259 for PI3P binding.

We also compared the activities of DenR WT vs. K259A using 17-ODYA in click chemistry assays. While robust formation of an acyl-enzyme intermediate (**Supporting Fig. 3A**) as well as substrate lipidation of GFP-NRAS (**Figure 3C**) and GFP-HRAS (**Supporting Fig. 3E**) was observed in cells producing mCherry-DenR, lysate from cells producing DenR(K259A) showed strongly reduced formation of a covalent reaction intermediate as well as lipidation of GFP-NRAS or -HRAS within transiently transfected HEK293 cells treated with 2BP, underscoring a critical role of PI3P recognition in regulating the catalytic activity of DenR.

### PI3P binding by the CTD allosterically activates the NTD

In all orthologs of DenR, the N-terminal NlpC/P60 domain is connected to the lipid-binding CTD via a linker displaying a proline (P)-rich patch (**Figure 1A**). Several residues in proximity were highly conserved among P4M-like homologs yet not predicted to directly contribute to either PI3P binding or catalysis, most notably F260 and Y267 within the CTD and P218 and P221 in the predicted flexible P-rich path of the linker region (**Figure 5F**). The Alphafold 3 structures suggested that this region adopts an isolated extended conformation forming a flexible tail that is likely to undergo a conformational change. Thus, it could serve as a conduit for a signal relay between the P4M-like CTD and the catalytic NTD where binding of the lipid head group to the P4M-like domain may allosterically regulate the activity of DenR. To investigate possible coordinated function between the CTD and NTD, we replaced several residues within the conserved linker region with alanine and performed click chemistry experiments to examine DenR activity. While P218 and P221 are highly conserved across all homologs, the Philadelphia-1 DenR sequence encodes five prolines in this region (**Supporting Fig. 5A**). DenR variants in which either two (P218A, P221A) or four (P218-221A) proline residues have been substituted with alanine (referred to as “APPA” and “AAAA” mutants, respectively) were created. Formation of an acyl-enzyme intermediate by DenR was significantly reduced with both linker mutations (**Supporting Fig. 5B**) while NRAS (**Figure 5G**, **Supporting Fig. 5C**) and HRAS (**Figure 5H**, **Supporting Fig. 5D**) substrate lipidation was significantly reduced with the DenR(AAAA) mutation. An allosteric activation model would also predict that disruption of PI3P binding to CTD would prevent S-acyltransferase activity by the NTD and consequently NRAS recruitment by DenR even if it was anchored to membranes via an alternative mechanism. To test this, we performed knock-sideways assays in which we equipped DenR(WT) or DenR(K259A) with the mitochondria targeting sequence MAO and assessed their ability to recruit GFP-NRAS in transiently transfected U2OS cells. DenR(WT)-mCherry-MAO efficiently redirected GFP-NRAS to the surface of mitochondria (**Figure 5I**). In contrast, DenR(K259A)-mCherry-MAO, despite being equally enriched on mitochondria, failed to recruit NRAS which remained localized to the PM and the Golgi. Similar results were obtained in HEK cells (**Supporting Fig. 5E**) where Pearson colocalization coefficients between DenR and NRAS signals revealed a statistically significant difference in NRAS recruitment between DenR wild-type and the PI3P-binding mutant (**Figure 5J**). In summary, these results support the hypothesis that PI3P binding by the CTD allosterically activates palmitoyltransferase activity of the NTD.

## Discussion

In this study, we demonstrate that *L. pneumophila* DenR is a bacterial S-acyltransferase that covalently modifies the proto-oncoprotein NRAS with palmitic acid in order to anchor it to LCV membranes during infection. This finding is significant for several reasons: First, to our knowledge, DenR is the first example of an S-palmitoyltransferase encoded by a bacterium.

Second, DenR challenges the conventional belief that palmitoyltransferases need to be integral membrane proteins and, instead, catalyzes the palmitoylation reaction while being exposed to the aqueous environment of the cytosol (**Supporting Fig. 6A**). Third, unlike DHHC enzymes, DenR is insensitive to the inhibitor 2BP (**Figures 2, 3**), suggesting a catalytic mechanism different from that of the DHHC proteins. Finally, palmitoyltransferase activity within the NTD required PI3P binding by the CTD of DenR and relay of the lipid binding signal through the linker domain (**Figure 5**), adding yet another example to the growing list of bacterial effectors that are regulated by allosteric activation. Circularly permuted versions of the NlpC/P60 clade of papain-like domains are found in a wide range of proteins from bacteria and archaea, but also eukaryotes (**Figure 1**). While several of these have been shown to operate on lipid/fatty acyl substrates, many of their actual activities in cellular contexts remain poorly understood. Thus, given the precedent established here, it is conceivable that some of these proteins may function as S-acyltransferases in taxa outside the Legionellaceae.

The atomic structures of two DHHC enzymes, human DHHC20 (hDHHC20) and zebrafish DHHC15 (zfDHHS15), recently revealed that although their transmembrane regions that form the acyl chain binding pocket are submerged in the hydrophobic core of the membrane, their catalytic D-H-H-C motif housed within the cysteine-rich domain was positioned outside the plane of the membrane (26). Upon placement of DenR onto a theoretical membrane (**Supporting Fig. 6A**), we found that the active site residues in DenR could be positioned similarly close to the lipid-water interface, likely via partial membrane insertion of the amphipathic helix, corresponding to the predicted cap region (**Figure 1D**) atop the active site, suggesting that the palmitoylation reaction can occur despite the bulk of DenR being exposed to the cytosol.

In DHHC enzymes, fatty acyl chain selectivity is determined by a hydrophobic cavity that is formed by the transmembrane domains and that accommodates fatty acid chains up to a certain length (26, 27). Although our experiments demonstrate the incorporation of the palmitic acid analog 17-ODYA into NRAS in a DenR-dependent manner, we cannot exclude the possibility that DenR transfers acyl chains other than palmitate to NRAS or that acyl donors other than palmitoyl-CoA serve as physiological substrates. Such promiscuity may be advantageous during infection of different host cell types (species of amoeba or macrophages), where the availability of palmitic acid within the LCV membrane may fluctuate over time. Thus, in order for DenR to efficiently recruit NRAS throughout the course of an infection, a constant supply of acyl chains of varying length that can be attached to NRAS may be more important than attaching acyl chains of a specific length, making a molecular measuring tape like the one described for DHHC enzyme unnecessary. Whether DenR can utilize substrates other than palmitoyl-CoA as acyl donors to catalyze the transfer of acyl groups to NRAS remains to be determined.

The finding that the CTD of DenR is a membrane targeting domain with specificity for PI3P (**Figure 5**) was unexpected, given that PI4P is the predominant phospholipid on membranes of the intermediate or late LCV when DenR is most active. We speculate that attraction towards the less abundant PI3P phospholipid might have evolved to prevent accidental spill-over of DenR to bystander organelles that are rich in PI4P, such as the Golgi, as this may negatively impact organelle function. Confining the activity of DenR to PI3P-positive organelles like the LCV at the expense of not being allowed to utilize the more abundant phospholipid PI4P may have proven to be advantageous. Although mitochondrial membranes contain relatively low levels of PI3P, residual PI3P may be sufficient for DenR to be activated when artificially redirected to this compartment and recruit GFP-NRAS, whereas the PI3P-binding mutant DenR(K259A) did not (**Figure 5I, J**). While the molecular mechanism underlying allosteric activation via different ligands remains to be better understood at the structural level, our findings provide important new insights into novel ways in which pathogens can exploit protein palmitoylation within eukaryotic cells without relying on host cell DHHC enzymes.

## Materials and Methods

### *L. pneumophila* strains and growth conditions

*L. pneumophila* strains were grown and maintained as described previously (28). Briefly, *L. pneumophila* cultures were grown overnight in ACES-buffered yeast extract and supplemented with <u>thymidine</u> (100 μg/mL), <u>chloramphenicol</u> (5 μg/mL), <u>kanamycin</u> (30 μg/mL), or IPTG (0.5 mM) as needed. The next day, cultures that had reached OD600 3–5 (post-exponential growth) were collected for use in host cell infections. All L. *pneumophila* cultures were visually inspected for appropriate morphology and motility prior to use. *L. pneumophila* strains Lp02 and Lp03 are thymidine-auxotroph derivatives of *L. pneumophila* strain Philadelphia-1 (29). All strains have been verified by whole genome sequencing.

### Plasmids

Plasmids used in this study were made using InFusion cloning technique according to the manufacturer protocol (Clontech) or using the Gibson cloning technique according to the manufacturer protocol (NEB), and mutant plasmids were made using site-directed mutagenesis (QuikChange Lightning Site-Directed Mutagenesis Kit, Agilent, #210519). pmCherry-C1 empty vector was purchased from Clontech (Cat#632524). Bacterial complementation plasmids encoding suntag-DenR(H25A) or DenR(H25A, E83A, C169A) in pJB908-mCherry vectors were custom ordered from GenScript. pmApple-SiT was a gift from Dr. Michael Davidson (Addgene plasmid #54948), GFP-NRAS and GFP-HRAS were a gift from Dr. Julie Donaldson, mCherry-Rab5A was a gift from Dr. Juan Bonifacino (30), mCherry-P4M-SidM was a gift from Dr. Tamas Balla (Addgene plasmid # 51471), and pmCherry-2xFYVE was a gift from Dr. Harald Stenmark (Addgene plasmid # 140050). mCherry-NRAS, GFP-DenR, mCherry-DenR, DenR-mCherry-MAO, pXDC50-mCherry, pJB908-mCherry, and pMMBC207C-DenR were generated as previously described (9). GFP-GobX 171-190 was generated as previously described (16). EBFP2-Rab7A was generated from GFP-canine Rab7A (a gift from Dr. Juan Bonifacino) by shuttling the Rab7A sequence into the EBFP2-C1 vector (a gift from Dr. Michael Davidson; Addgene #54665) using KpnI-HF and ApaI digestion of both donor and acceptor plasmids. pGST-parallel1 was a gift from Dr. Juan Bonifacino. GST-SidM was a gift from Dr.

Romona Neunuebel (31).

DenR-K259A-mCherry-MAO was generated using QuikChange site-directed mutagenesis of DenR-mCherry-MAO using forward (5’ – GTCCCATGACCAAGCGCGCATTTAATAGCCTCAAGAACCTGTATCTCC) and reverse (5’ - GGAGATACAGGTTCTTGAGGCTATTAAATGCGCGCTTGGTCATGGGAC) oligos.

GFP-HRAS(C184S) was generated using QuikChange site-directed mutagenesis of GFP-HRAS using forward (5’ – GAGTGGCCCCGGCTGCATGGCACTCAAGTGTGTGCTCTCCTGAGGGCCCGGG) and reverse (5’ – CCCGGGCCCTCAGGAGAGCACACACTTGAGTGCCATGCAGCCGGGGCCACTC) oligos.

GFP-DenR(CTD) and GFP-DenR(NTD) were generated using QuikChange site-directed mutagenesis of GFP-DenR using the following oligos (**for CTD:** forward 5’ – CGAGCTCAAGCTTCGGCCACCCCGCTGGTACAGAACGGCAACGC, reverse 5’ – GCGTTGCCGTTCTGTACCAGCGGGGTGGCCGAAGCTTGAGCTCG; **for NTD:** forward 5’ – CCATTCTATGTTCTCCCGCCACCTTGTCCCTACTAGTCGACGGTACCGC, reverse 5’ - GCGGTACCGTCGACTAGTAGGGACAAGGTGGCGGGAGAACATAGAATGG).

A codon optimized gBlock encoding Lmic2405(CTD) (5’ - CTCAGATCTCGAGCTCAAGCTTCGTTTTACGTCTTTCCTCTGCCTCCAGACGGATACCCAGC AGACGAAACAAAAAAGGCAATACTCACAAAAATCTTCCGCCGGATGGAAAATCTCTTGAAAA AAGATCCCTATAGCGAGAATACAATCCAGAAGTTTGAAGCCCTTAAAAACCTTTACAACGAG CAAGCTGGGATCCCTGATAATGACATAGAGCAGGCGTTGAAATCCATTCGGATATGGAAATC TACCAACCACGATGTCATTTGTCAGTTGCGAGCCCAATCTTTCCTCGGGAAATTGTTCGTCA GCACCTCATCCACGAAGGCTATGGCCGATGACATAGAGGATGCACTGGAGCGGGACCTGG TACTTtagcaGTCGACGGTACCGCGGGCCCG) was purchased from IDT, digested with BglII and SalI restrictions enzymes and ligated into pmCherry-C1 double digested with the same enzymes to generate mCherry-Lmic(CTD). The BglII and SalI double digested gBlock was also ligated into pGST-parallel1 vector digested with BamHI and SalI. The resulting construct was then mutated to correct its reading frame using QuikChange site-directed mutagenesis and forward (5’ – CCTGTATTTTCAGGGCGCCATGAGATCTCGAGCTCAAGCTTCG) and reverse (5’ – CGAAGCTTGAGCTCGAGATCTCATGGCGCCCTGAAAATACAGG) oligos. GST-Lmic(CTD) was the final, frame corrected, construct.

Plasmids encoding DenR with point mutations were generated using QuikChange site-directed mutagenesis of either GFP- or mCherry-tagged DenR plasmids. For GFP-DenR(CTD-K259A) and GST-DenR(CTD-K259A), forward (5’ – GTCCCATGACCAAGCGCGCATTTAATAGCCTCAAGAACCTGTATCTCC) and reverse (5’ – GGAGATACAGGTTCTTGAGGCTATTAAATGCGCGCTTGGTCATGGGAC) oligos were used.

For mCherry-DenR(H25A), forward (5’ - CAGCATAGGAAAAATTAACCAAGCGTCATTTATCATGCTCGGCG) and reverse (5’ - CGCCGAGCATGATAAATGACGCTTGGTTAATTTTTCCTATGCTG) oligos were used. For mCherry-DenR(E83A), forward (5’ - GGCAAAGATTGAAGATGCAGGTATTACTCGCAAAAGAAAGGGG) and reverse (5’ - CCCCTTTCTTTTGCGAGTAATACCTGCATCTTCAATCTTTGCC) oligos were used. For mCherry-DenR(C169A), forward (5’ - CACAGGAAATACCGCGAGACATTCTGCTATCAAGC) and reverse (5’ - GCTTGATAGCAGAATGTCTCGCGGTATTTCCTGTG) oligos were used. For mCherry-DenR(L68A), forward (5’ - GTCTCTCAATCTACTCTGCCGCCAGTGGTGCTATTTTTGCC) and reverse (5’ - GGCAAAAATAGCACCACTGGCGGCAGAGTAGATTGAGAGAC) oligos were used. For mCherry-DenR(APPA), forward (5’ - CATTCCATTCTATGTTCTCGCGCCACCTGCACCGAGCATACATCCGGG) and reverse (5’- CCCGGATGTATGCTCGGTGCAGGTGGCGCGAGAACATAGAATGGAATG) oligos were used.

For mCherry-DenR(AAAA), forward (5’ - CTCTCAAAACATTCCATTCTATGTTCTCGCGGCAGCTGCACCGAGCATACATCCGGG) and reverse (5’ - CCCGGATGTATGCTCGGTGCAGCTGCCGCGAGAACATAGAATGGAATGTTTTGAGAG) oligos were used.

To generate GST-DenR(CTD), the DenR(CTD) domain was digested from GFP-DenR(CTD) with SacI and SalI enzymes and ligated to pGST-parallel1 vector digested with SacI and XhoI enzymes. For full length GST-DenR constructs, GST-DenR(CTD) was double digested with HindIII-HF and XhoI and the backbone vector gel purified. mCherry-DenR(WT), mCherry-DenR(H25A), mCherry-DenR(E83A), mCherry-DenR(C169A), and mCherry-DenR(K259A), were each double digested with HindIII-HF and SalI-HF and DenR sequences were gel purified. These DenR sequences were ligated to the purified vector backbone to create corresponding GST-tagged constructs.

### Cell lines

HT1080 cells (ATCC CCL-121), HeLa cells (ATCC CCL-2), HEK293T cells (ATCC CRL-3216), and U2OS cells (ATCC HTB-96) were obtained from the American Type Culture Collection (ATCC) and verified to be mycoplasma free. An HT1080 cell line stably producing GFP-NRAS was previously created, as described (9), and used for infection assays in **Figures 2C, 2D**, and **Supporting Fig. 2C**. All cell lines were maintained in Dulbecco’s Modified Eagle Medium containing 1.5 g/L sodium bicarbonate and L-glutamine (Quality Biological; # 112-319-101) supplemented with 10% fetal bovine serum (FBS; Corning, #35-011-CV) and 100 IU/mL penicillin and 100 μg/mL streptomycin (Corning, #30-002-CI) (CDMEM) in humidified incubators at 37°C and under 5% CO_2_. Cell lines were routinely examined for absence of mycoplasma contamination by assessing cytoplasmic DAPI staining of fixed samples of cell cultures.

### In silico analysis

#### Alphafold modeling

The amino acid sequence of DenR (Lpg1909) from *Legionella pneumophila* was entered into Alphafold 2 (v2.3.2, NIH HPC Biowulf cluster) and Alphafold 3 (https://alphafoldserver.com/) to predict its domain architecture. Both programs predicted similar domain architecture. Alphafold 2 was used to generate multiple sequence alignments (MSAs), while Alphafold 3 was used for monomer and multimer modelling. The Alphafold 3 DenR monomer prediction had a confidence score of pTM=0.88. For modeling the heterodimer shown in **Figure 2A**, sequences for NRAS and DenR (lpg1909) were entered into Alphafold 3. The V187/V188/M189 residues were deleted from the C-terminal NRAS sequence to mimic CAAX protease processing, and a palmitoylation moiety was added to C186 to mimic protein acylation at this site (farnesylation was not an available modification choice in the Alphafold 3 online server) prior to running model prediction. A palmitic acid ligand was also added for the model prediction. All predicted models were viewed, manipulated, and rendered with ChimeraX v1.7.1.

#### DenR domain analysis

ChimeraX was used to create a PDB file of the DenR (Lpg1909) monomer. This was uploaded to the Dali protein structure comparison server (http://ekhidna2.biocenter.helsinki.fi/dali/) to identify proteins with predicted functional homology based on structure (**Figure 1A**).

#### DenR homolog comparison

Multiple sequence alignments (MSAs) generated by Alphafold 2 were inspected for unique protein identifiers (NCBI or Uniprot) using Python, recovering 98 discrete identifiers from 48 different species of *Legionella* bacteria plus 9 species which were unspecified. Protein hits from the 9 unspecified *Legionella* were discounted. A single protein sequence from each of the remaining *Legionella* species was chosen as a DenR homolog representative for that species (48 total). When a single species contained multiple predicted homologs, those with an identified gene name [e.g. Lpar2301 or Lmic2405] were given preference, otherwise those with best sequence conservation to DenR were chosen. These 48 DenR homologs were sequence aligned using Clustal Omega(1.2.4) on the HHPRED server (https://toolkit.tuebingen.mpg.de/tools/hhpred). The DenR lpg1909 Alphafold 3 monomer model and Clustal O sequence alignments (.clustal file) were imported together into ChimeraX. A DenR sequence conservation model is presented in **Supporting Fig. 1C** as the Lgp1909 monomer model color-coded at each residue with a sequence conservation score, using the “structural analysis > render by attribute” feature in ChimeraX.

Alphafold 3 monomer predictions were generated of all 48 DenR homologs and imported together into ChimeraX. Using the “structure analysis > render by attribute” feature, root mean squared displacement (RSMD) values of imported models along each residue of lpg1909 were calculated. The structural homology of DenR homologs is presented in **Supporting Fig. 1D** with the lpg1909 monomer model color-coded by these RSMD values.

Profile-Profile searches were run against profiles built from the Pfam and PDB databases using the HHpred program. Domains were then identified by deploying a collection of HMMs and PSSMs maintained by the Aravind lab, along with Pfam HMMs, using RPSBLAST and HMMSCAN. To further refine detection, domain identification was extended through remote homology analysis using HHpred against profiles built from the Pfam and PDB databases.

Phylogenetic analysis of the NlpC/P60 domains was performed with FastTree and rendered with TreeViewer. The domain architectures were compiled from the significant matches in the above-described profile searches and superimposed onto branches of the trees. The branches were colored according to their prevalent contextual connections and rooted based on the conservation pattern of the active motif of the NlpC/P60 domain. The multiple sequence alignment was used to compute bit-score based sequence logos for each of the families depicted in the tree.

### Plasmid DNA transfection

Plasmids were delivered into cells using Lipofectamine 2000 (Life Technologies, # 11668019) according to the manufacturer protocol. For live-cell imaging, cells were seeded to 4- or 8-well chambered cover glass (Cellvis #C4-1.5H-N or #C8-1.5H-N), coated with collagen (Gibco #A10644-01) according to the manufacturer protocol. After 24 h, cells were transfected with plasmid DNA. Transfection complexes were prepared in Opti-MEM Reduced Serum Medium (Thermo Fisher Scientific, #31985062), using the equivalent of 0.1 μg of each plasmid and 0.5 μL Lipofectamine 2000 per well of an 8-well chamber, and overlaid onto cells. Culture media was replaced by fresh CDMEM 3-6 h after transfection. 16-24 h later, cells were either imaged by live microscopy or were challenged with Lp and imaged live 12 hours after infection (**Figure 4C**).

### Live confocal microscopy

For live-cell imaging, cells were seeded to 4- or 8-well chambered cover glass (Cellvis #C4-1.5H-N or #C8-1.5H-N), coated with collagen (Gibco #A10644-01) according to the manufacturer protocol. All live-cell imaging was performed on either a Zeiss LSM780, LSM800, or LSM880 confocal laser scanning microscope equipped with an incubated chamber (5% CO_2_ and 37°C) and a Plan-Apochromat 63X, 1.4 numerical aperture (NA) objective (Carl Zeiss). Microscopy images were acquired with Zeiss ZEN Black software (Zen 2.3 SP1 FP3 release version 14.0.25.201) or ZEN Blue software (Zen 3.9 version 3.9.101.11000). Image processing, intensity measurements, and quantification were performed using Fiji v1.54p (NIH, Bethesda, MD). All scale bars drawn to 5 μm unless otherwise stated.

For experiments involving host cell challenge with Lp (**Figures 2C, 2D, 4C**, and **Supporting Fig. 2C**), HT1080 cells plated on collagen-coated, chambered coverglass were cultured for at least 1 h with complete media lacking antibiotics (infection media) prior to challenge. Bacterial strains were diluted in infection media and added to HT1080 cells at an MOI 25–50. In all cases, bacteria were centrifuged onto the cells for 5 min at 200 *x g* prior to incubation. After 1 h of incubation, cell monolayers were washed three times with PBS to remove extracellular bacteria and replaced with fresh infection media. Cells were then incubated for the indicated times prior to imaging by live microscopy. The quantification of NRAS localization to LCVs in HT1080 cells (**Figure 2C and Supporting Fig. 2C**) was performed by imaging random fields of live infected cells by confocal microscopy and counting the percentage of infected cells displaying distinct NRAS signal along the full limiting membrane of the LCV. HT1080 cells seeded to collagen-coated coverglass were challenged with Lp02Δ*denR* complemented with DenR(WT), suntag-DenR(H25A), or DenR(HEC) in pJB908-mCherry vectors at MOI 25 for the indicated times (either 6-8 h or 16-18h) prior to live-cell imaging. Data are presented as the mean ± SD from scoring 50-66 LCVs per condition for three independent experiments. Statistical significance was calculated using one-way ANOVA with Tukey’s multiple comparisons test to determine an adjusted p-value for each comparison.

The quantification of GFP-NRAS recruitment to mCherry-DenR-positive intracellular membranes in **Figure2B** was performed by live-cell imaging transiently transfected HT1080 cells seeded on collagen-coated coverglass. Population counts of transfected cells displaying obvious NRAS localization to intracellular membranes positive for mCherry-DenR are presented as the mean ± SD from scoring 40-80 co-transfected cells for each condition from three independent experiments.

The quantification of GFP-HRAS enrichment on mCherry-DenR positive endomembranes in the presence of 2BP in transiently transfected HeLa cells (**Figure 3E**) was performed by imaging random fields of cells by confocal microscopy and performing maximum intensity projections of GFP-HRAS signals from collected z-stacks for each condition. Images from a single representative experiment were thresholded to exclude uniform, low-intensity signal; leaving bright signal intensities associated with endomembrane recruitment. Fluorescence intensity values above threshold per cell were recorded in at least 30 cells from each condition and presented as mean ± SD, with individual dots representing single cells. Quantification of GFP-HRAS(C184S) to mCherry-DenR positive endomembranes in the presence of 2BP in transiently transfected HT1080 cells (**Supporting Fig. 3B**) was performed by imaging random fields of live cells by confocal microscopy and counting the percentage of co-transfected cells with GFP-HRAS(C184S) signal readily observable as bright endomembrane puncta. Data are presented as mean ± SD from scoring 50-100 co-transfected cells per condition in two independent experiments.

To quantify GFP-NRAS recruitment to mitochondria membranes (**Figures 2E and Supporting Fig. 2E**), HT1080 cells were transiently transfected to produce GFP-NRAS with either DenR-mCherry-MAO or dsRed1-mito. The following day, cells were treated with 75μM 2BP and immediately imaged live at 20 sec intervals for 49 min (**Figure 2E**) or 66 min (**Supporting Fig. 2E**). Still frame images of GFP-NRAS and a mitochondrial marker (DenR-mCherry-MAO or dsRed1-mito) at the beginning and end of the acute 2BP treatment are displayed in the figures using the ImageJ/Fiji ICA colormap to enhance contrast of membrane-associated signals.

Automatic thresholding of the mitochondrial marker signal (either DenR-mCherry-MAO or dsRed1-mito) was used to create a mask of mitochondrial membranes for each acquisition frame. Signal intensities in each channel at these regions were then quantified and plotted for the representative cell to show mitochondria-specific signal.

### Quantification and statistical analysis

Statistical analyses were performed using GraphPad Prism 10 software (version 10.5.0), applying two-tailed, unpaired Student’s t-test when two groups were compared or one-way ANOVA analysis when multiple groups were compared. p values <0.05 were considered as statistically significant.

### GST-fusion bead generation

BL21-Gold (DE3)pLysS (Agilent #230134) bacteria were transformed with indicated plasmids for either protein purification in the PIP strip assay (GST-Par1 [NovoPro #V005870], GST-SidM, GST-DenR(CTD), GST-DenR(CTD-K259A), or GST-Lmic(CTD)) or GST-DenR(K259A)), or to make mCherry (pGEX6P1-mCherry-nanobody [Addgene #70696]) and GFP (pGEX6P1-GFP-nanobody [Addgene #61838]) nanobody beads for the Click Chemistry assays. Transformed bacteria were grown on LB+Amp selection plates. Individual colonies were inoculated into a starter culture of 5 mL LB supplemented with 100 μg/mL ampicillin, and grown overnight at 37 °C with moderate shaking. The entire 5 mL inoculum was transferred to 250 mL LB supplemented with 100 μg/mL ampicillin and 1mM IPTG and grown overnight at 19°C with moderate shaking.

Bacteria were then lysed by sonication in lysis buffer (50mM Tris [pH8.0], 150 mM NaCl, 5 mM DTT, 100 μg/mL DNAse, 100 μg/mL lysozyme, complete-EDTA-free protease inhibitor [Roche #1183617001]) and centrifuged for 30 min at 14,000 x g at 4°C. The resulting supernatant was collected and incubated with 400 μL glutathione Sepharose beads (Cytiva #17075601) at 4°C with continuous rotation. Beads were pelleted by centrifugation at 500 x g for 1min at 4°C, resuspended in bead wash buffer (50mM Tris [pH 8.0], 150mM NaCl, 5mM DTT, complete-EDTA-free protease inhibitor), and stored at 4°C.

### GST-DenR pulldown assays

GST-DenR(WT), GST-DenR(H25A), GST-DenR(E83A), GST-DenR(C169A), or GST-DenR(K259A) were expressed in BL21-Gold (DE3)pLysS (Agilent #230134) bacteria as above and induced with 1 mM IPTG for 5 h at 37°C with moderate shaking. Bacterial cells were harvested and lysed in buffer containing 1× PBS, 5% glycerol, 1% CHAPS, and protease inhibitors. Lysates were clarified by centrifugation and GST–DenR fusion proteins were immobilized on GST affinity beads. The beads were washed with 30 bead volumes of 1× PBS, 5% glycerol, 300mM NaCl, and the bead-bound proteins were retained for subsequent pull-down assays.

GFP-tagged NRAS was expressed in HEK 293T cells. Cells were lysed in buffer containing 20 mM HEPES, pH 7.2, 150 mM NaCl, 1% NP-40, and protease inhibitors. Bead-bound GST–DenR proteins were incubated with the GFP–NRAS-expressing cell lysates for 4 h at 4°C. Following incubation, the beads were washed twice with buffer containing 20 mM HEPES, pH 7.2, 300 mM NaCl, and 1% NP-40. Bound proteins were recovered by adding 1× Laemmli sample buffer and heating at 90°C for 10 min. Samples were separated by SDS–PAGE and analyzed by immunoblotting with rabbit anti-GST antibody (Thermo Fisher #MA541079) and rabbit anti-GFP antibodies (GenScript #A01388-40) to detect GST–DenR bait proteins and associated GFP–NRAS, respectively.

### PIP strip assay

GST-beads (GST-Par1, GST-SidM, GST-DenR(CTD), GST-DenR(CTD) K259A, or GST-Lmic CTD) generated as described above, were washed three times in bead wash buffer (50mM Tris [pH 8.0], 150mM NaCl, 5mM DTT, complete-EDTA-free protease inhibitor) and bound proteins were eluted from beads in 400 μL bead elution buffer (50 mM Tris [pH 8.0], 10 mM reduced glutathione (Sigma G4251-25G), complete-EDTA-free protease inhibitor). 2.5 μg of protein eluate was diluted in 5 mL PIP strip blocking buffer (10mM Tris [pH 8.0], 150mM NaCl, 0.1% Tween20, 3% FA-free BSA [Sigma #A7030-100G]) and overlaid onto a PIP strip membrane (Invitrogen #P23751) that had previously been incubated in blocking buffer overnight at 4°C with gentle shaking. The PIP strip membranes were incubated in the protein solution overnight at 4°C with gentle shaking, and was subsequently washed 3 times in PIP strip wash buffer (10 mM Tris [pH8.0], 150 mM NaCl, 0.1% Tween20). Goat anti-GST antibody (Sigma #GE27-4577-01) was then diluted 1:5,000 in PIP strip blocking buffer and incubated on the membranes for 1 h at RT with gentle agitation. Membranes were washed again 3 times in PIP strip wash buffer, followed by 1 h incubation at RT with donkey anti-goat HRP secondary antibody diluted 1:5,000 in PIP strip blocking buffer. Membranes were again washed 3 times in PIP strip wash buffer, developed with SuperSignal West Dura Extended Duration Substrate (Thermo Fisher #34075), and signal detected using a Bio-Rad ChemiDoc MP imaging system.

### Acyl-Resin Assisted Capture (Acyl-RAC) assay

Acyl-RAC procedures were carried out similar to a recent Bio-protocol publication (32). Briefly, 2.5 x 10^6 HEK293T cells were seeded to 10-cm dishes. After 24 h, cells were transfected with plasmid encoding either GFP-NRAS alone, or GFP-NRAS and mCherry-DenR, using Lipofectamine 2000. Transfection complexes were prepared in Opti-MEM Reduced Serum, using the equivalent of 4 μg of each plasmid and 12.5 μL Lipofectamine 2000 for each 10-cm dish, and overlaid onto cells. Culture media was replaced by fresh CDMEM supplemented with 2BP (25 μM), 3-6 h after transfection, and cells incubated overnight. The following day, 1 μM palmostatin B (PalmB; Sigma #508738) was added to media to inhibit protein depalmitoylation and incubated with cells for 1 h at 37°C under 5% CO_2_. Cells were lifted from the dish with cell lifter (Corning #3008), collected with the media, and pelleted by centrifugation at 300 x g for 5 min at 4°C. Pellets were washed once in PBS supplemented with 1 μM PalmB, followed by resuspension in 500 μL Acyl-RAC lysis buffer (25 mM HEPES, 25 mM NaCl, 1 mM EDTA, 1.5% Triton X-100, 1 μM PalmB, and complete-EDTA-free protease inhibitor, pH 7.5). Resuspended pellets were incubated for 1 h at 4°C with continuous rotation. Lysates were then centrifuged at 16,000 x g for 15 min at 4°C, and SN collected. TCEP was added to each SN to a final concentration of 50 mM, and samples incubated at RT for 1h with continuous rotation to reduce disulfide bonds in non-acylated proteins. 1 mL Acyl-RAC blocking buffer (100 mM HEPES, 1 mM EDTA, 2.5% SDS, 1.5% MMTS, pH 7.5) was then added to each and samples were incubated for 4 h at 40°C with gentle agitation. 6 mL of ice-cold acetone were added to each sample, vortexed, and incubated for 48 h at −20°C. Samples were centrifuged at 5,000 x g for 10 min at 4°C and pellets washed 4 times with 1 mL 70% acetone, then air-dried. Dried pellets were resuspended in 500 μL Acyl-RAC binding buffer (100 mM HEPES, 1 mM EDTA, 1% SDS, pH 7.5), rotated at RT until completely dissolved, and then divided into two aliquots and supplemented with 100 μL of either 2 M NaCl (negative control) or freshly prepared 2 M hydroxylamine (pH 7.4) (to specifically cleave thioester bonds). 50 μL Thiopropyl Sepharose beads (G-Biosciences #786-1785) per aliquot were washed with distilled water, centrifuged at 2,500 x g for 10min, resuspended in Acyl-RAC binding buffer, and incubated overnight at RT with continuous rotation and protected from light. The following day beads were washed 5 times with Acyl-RAC binding buffer, being centrifuged at 9,000 x g for 5 min at RT after each wash, resuspended in 4X reducing SDS-PAGE buffer, and heated at 95 for 10 min. Beads were then centrifuged at 2,500 x g for 5 min and eluates (SN) collected. Equal sample volumes of cell lysates or final eluates were then subjected to immunoblotting.

### Immunoblots and antibodies

Cell lysates or samples as described above were incubated with sample buffer, boiled for 10 min, loaded on 4–12% Bis-Tris gels (Nu-PAGE, Invitrogen), and then transferred onto nitrocellulose membranes (GE Healthcare). The membranes were blocked in PBS (with 0.1% Tween-20, 3% non-fat-dried milk) for 1 h and incubated with respective primary and secondary antibodies diluted in blocking buffer for 1 h at RT. Membranes were washed 3 times with blocking buffer for 5 min each before and after secondary antibody incubation, developed with SuperSignal West Dura Extended Duration Substrate (Thermo Fisher #34075), and signal detected using a Bio-Rad ChemiDoc MP imaging system.

### Click chemistry

Click chemistry procedures were carried out similar to a recent protocol outline in JoVE (33). Briefly, 2.5 x 10^5 HEK cells were seeded to each well of a 6-well tissue culture plate. After 24 h, cells were transfected with plasmid encoding either GFP-NRAS or GFP-HRAS along with mCherry-DenR(WT), mCherry-DenR(CTD), or mCherry-DenR mutant constructs, using Lipofectamine 2000. Transfection complexes were prepared in Opti-MEM Reduced Serum, using the equivalent of 0.68 μg of each plasmid and 4 μL Lipofectamine 2000 for each well, and overlaid onto cells. Culture media was replaced by fresh CDMEM supplemented with 2BP (25 μM), 3-6 h after transfection, and cells incubated overnight. The following day, media was aspirated from cells and replaced with 2 mL pre-warmed DMEM supplemented with 5% charcoal stripped fetal bovine serum (Invitrogen #12676029) and 2BP (25 μM) for 1 hr. Fresh 17-ODYA was prepared by mixing in a glass vial 20 μL each of 100 mM 17-ODYA and 120 mM KOH for each well of the 6-well plate. The final mixture was added to cells (40 μL total per well) and incubated for 3-5 h at 37°C under 5% CO_2_. Media was then removed and cells were lifted from the dish with cell lifter (Corning #3008) in PBS supplemented with 1 μM palmostatin B. Lifted cells were pelleted by centrifugation at 300 x g for 5 min, and pellets incubated in 800 μL lysis buffer (25 mM HEPES pH7.4 + 0.5% IGEPAL-CA630 + 25 mM KCl + 2 mM MgCl2 + complete-EDTA-free protease inhibitor + 1 μM palmostatin B) at 4°C for 30 min, with continuous rotation. Lysates were centrifuged at 15,000 x g for 15 min at 4°C and 20 μL of either mCherry (autoacylation assays) or GFP (substrate acylation assays) nanobody beads, created as described above (in GST-fusion bead generation), were added to the SN and incubated overnight at 4°C with continuous rotation. The following day, beads were pelleted by centrifugation at 500 x g for 1 min, washed 3 times in wash buffer (25 mM HEPES pH7.4 + 0.05% IGEPAL-CA630 + 25 mM KCl + 2 mM MgCl2) and bound proteins eluted in 45 μL elution buffer (50 mM HEPES + 1% SDS) incubated at 80°C for 15 min with gentle agitation. To the SN, freshly made click chemistry mixture was added to a final concentration of 100 μM TAMRA, 100 μM TBTA, 1 mM CuSO4, and 1 mM TCEP. Click chemistry mixture was incubated at 37°C for 1 h with gentle agitation, then the reaction stopped by adding 4X SDS supplemented with 18% β-ME. This solution was incubated at RT for 10 min with gentle agitation and 20 μL each sample then immediately loaded to 4-12% Bis-Tris polyacrylamide gels. Gels were run until the dye front containing unbound TAMRA was completely run off the gel, to reduce background. TAMRA signal in the gels was then detected using a Bio-Rad ChemiDoc MP imaging system.

## Acknowledgments

This research was supported by the Intramural Research Program of the National Institutes of Health (1ZIAHD008893-16 to MPM). The contributions of the NIH author(s) are considered Works of the United States Government. The findings and conclusions presented in this paper are those of the author(s) and do not necessarily reflect the views of the NIH or the U.S. Department of Health and Human Services. The authors would like to thank Dr. Jennifer Kunselman for the generation of the pEBFP2-Rab7A plasmid, and Dominic Esposito, Daniel Bonsor, and Dhirendra Simanshu (RAS Initiative, Frederick National Laboratory for Cancer Research), and all members of the Machner lab for insightful discussions. This work utilized the computational resources of the NIH HPC Biowulf cluster (https://hpc.nih.gov).

## Author Contributions

C.D.W., S.B., and M.P.M. conceptualized the study; C.D.W., S.B., R.P., S.S.L., and G.G.N. performed the experiments; C.D.W., L.A., and M.P.M. wrote the manuscript; All authors analyzed the data, provided intellectual input into the study and contributed to editing of the manuscript; M.P.M. supervised and managed the entire project.

---

**Supporting Fig. 1:**
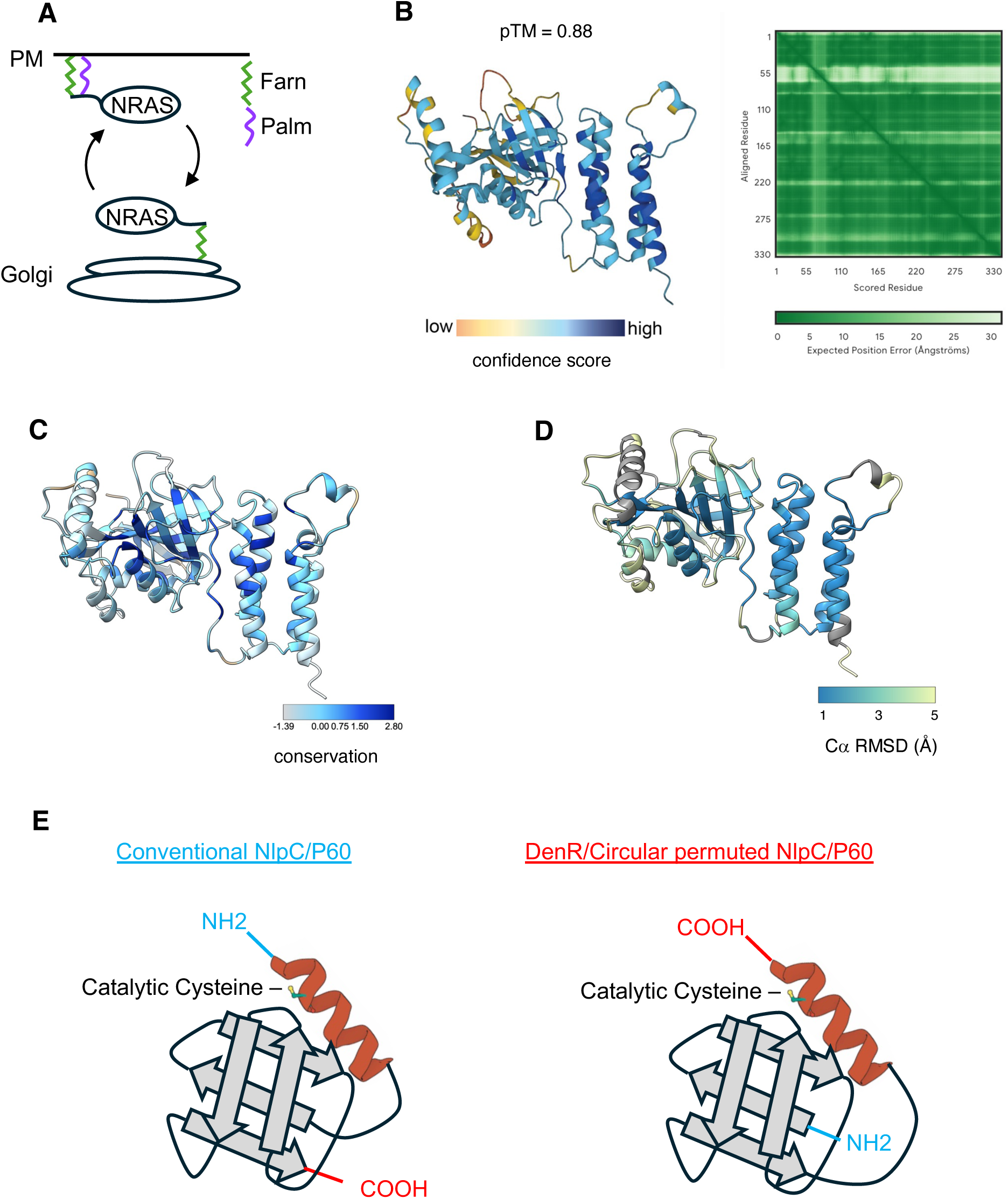
**(A)** Cartoon of NRAS membrane cycling. Palmitoylated NRAS localizes to the plasma membrane (PM), while depalmitoylated NRAS returns to the Golgi for another cycle of palmitoylation. **(B)** Alphafold 3 model showing confidence score as heat map along the peptide backbone of DenR. **(C)** Sequence conservation model of DenR homologs color-coded from light (least conserved) to dark (most conserved) at each alpha-carbon along the peptide backbone. **(D)** Structural homology model of DenR homologs color-coded by RMSD value at each alpha-carbon along the primary sequence. Regions of strongest structural conservation are colored in blue. **(E)** Model highlighting the different location of the helix containing the catalytic cysteine residue in conventional (left) vs circular permuted (right) NlpC/P60 domains.

**Supporting Fig. 2:**
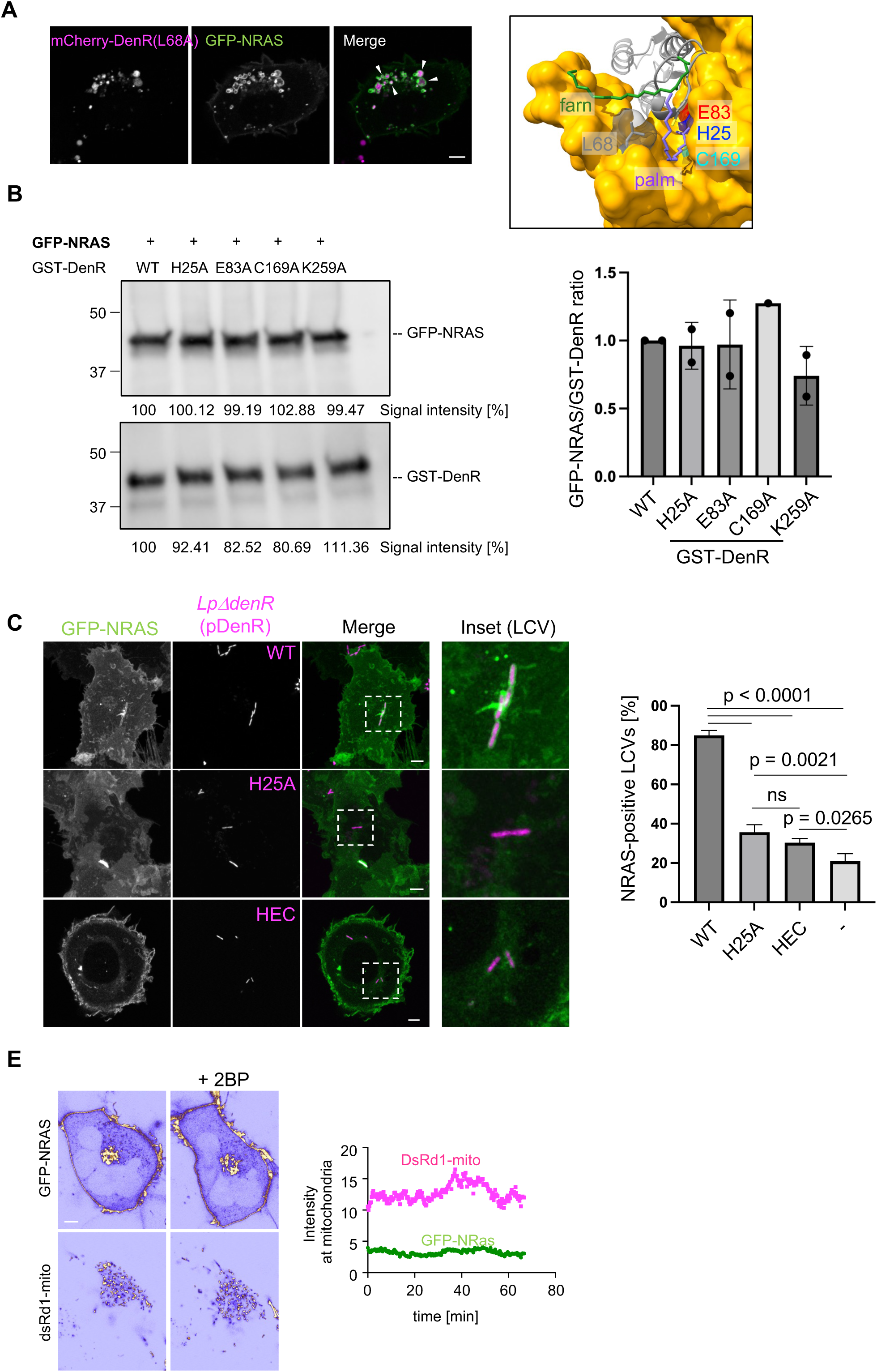
**(A)** DenR(L68A) mutant recruits NRAS to endomembranes. Transiently transfected HT1080 cells producing GFP-NRAS and mCherry-DenR(L68A) were imaged by fluorescence microscopy. Arrowheads indicate areas with NRAS recruitment to DenR(L68A)-positive membranes. Structural model shows the location of L68 relative to the active site. **(B)** Active site mutants of DenR retain substrate binding ability. GST-DenR wild-type or mutant proteins were purified from BL21 *Escherichia coli* using GST affinity beads and incubated with cell lysates from HEK cells producing GFP-NRAS. Pulldown assays tested the ability of GST-DenR to interact with GFP-NRAS substrate proteins. To the left, anti-GFP (top) and anti-GST (bottom) blots indicate that wild-type and mutant DenR proteins are similarly able to interact with NRAS. Graph to the right represents mean +/− SD quantification of the ratio of GFP-NRAS signal normalized to GST-DenR signal for each from two experiments (n=2), except for C169A which is from a single experiment (n=1). **(C)** Active site mutants of DenR fail to recruit NRAS to LCVs at 6 hpi. HT1080 cells producing GFP-NRAS were challenged with Lp02Δ*denR* complemented with plasmids encoding the indicated DenR variants, and GFP-NRAS colocalization with LCV was examined 6 hpi by fluorescence microscopy. Insets show magnified views of selected LCV displaying the prominent phenotype for each complement construct. Graph shows mean +/− SD quantification of three experiments (n=3). **(D)** Inhibition of cellular DHHC enzymes does not cause NRAS localization to mitochondrial membranes in the absence of DenR. As a control for Figure 3C, transiently transfected HT1080 cells producing dsRed1-mito and GFP-NRAS were treated with 2BP (75 μM), and recruitment of NRAS to mitochondria was examined by live fluorescence microscopy for 60 min. Micrographs from the start and end of the imaging series are displayed, pseudocolored using the ICA lookup table to better highlight changes in intensity. The graph quantifies NRAS and dsRed1-mito fluorescence intensity at mitochondria over time upon 2BP treatment in the representative cell displayed. All scale bars, 5 µm.

**Supporting Fig. 3:**
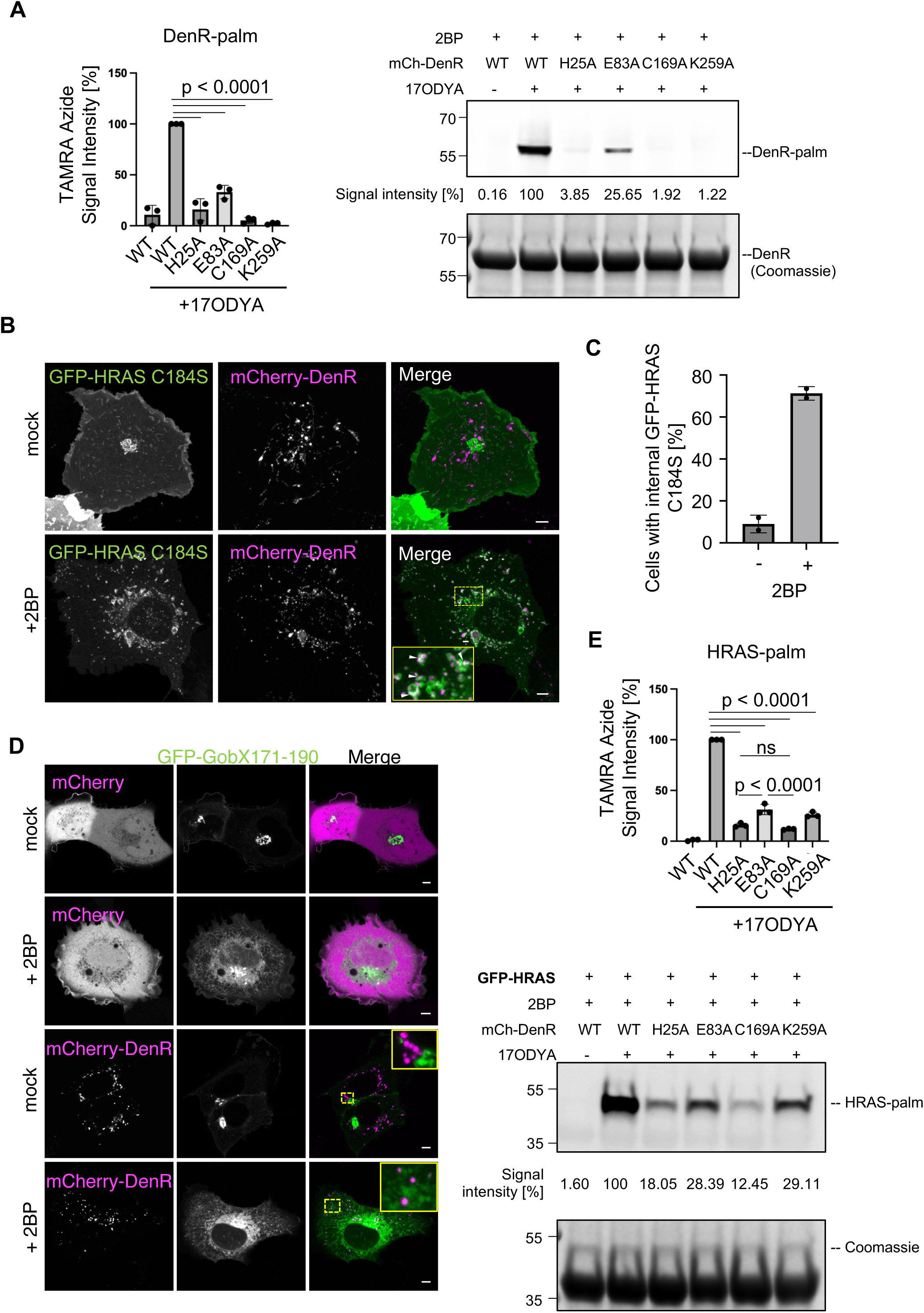
**(A)** Click chemistry to detect DenR autoacylation. 2BP-treated HEK293 cells producing the indicated mCherry-tagged DenR variants were metabolically labelled with 17ODYA. mCherry-tagged DenR was affinity-purified by immunoprecipitation with mCherry nanobody beads, and palmitoylation was detected by click chemistry with TAMRA-azide and in-gel visualization using a ChemiDoc Imaging System. Graph to left displays mean +/− SD for three experiments (n=3), while blots to right show a single representative experiment. Numbers indicate signal intensity relative to WT DenR with 17ODYA which was arbitrarily set to 100%. **(B)** Inhibition of cellular DHHC enzymes stimulates HRAS(C184S) recruitment by DenR. Transiently transfected HT1080 cells producing GFP-HRAS(C184S) and mCherry-DenR were treated with 2BP, and colocalization of NRAS with DenR on endomembranes was examined by fluorescence microscopy. Insets represent examples of endomembranes; white arrowheads indicate regions of colocalization. **(C)** Quantification of the percentage of the cellular population (mean +/− SD) exhibiting GFP-HRAS C184S recruitment to endomembranes (seen in A). 50-100 co-transfected cells counted per condition in two experiments (n=2). **(D)** The *L. pneumophila* effector GobX is not a substrate of DenR. HT1080 cells producing GFP-GobX171-190 and either mCherry or mCherry-DenR were mock-treated or treated with 2BP, and localization of both proteins was examined by fluorescence microscopy. Insets represent examples of endomembranes. **(E)** Same experiment as Figure 3C but with GFP-HRAS. Graph represents quantification of three experiments (n=3), mean +/− SD. Blots from a single representative experiment are shown below. Numbers indicate signal intensity relative to WT DenR with 17-ODYA which was arbitrarily set to 100%. All scale bars, 5 µm.

**Supporting Fig. 4:**
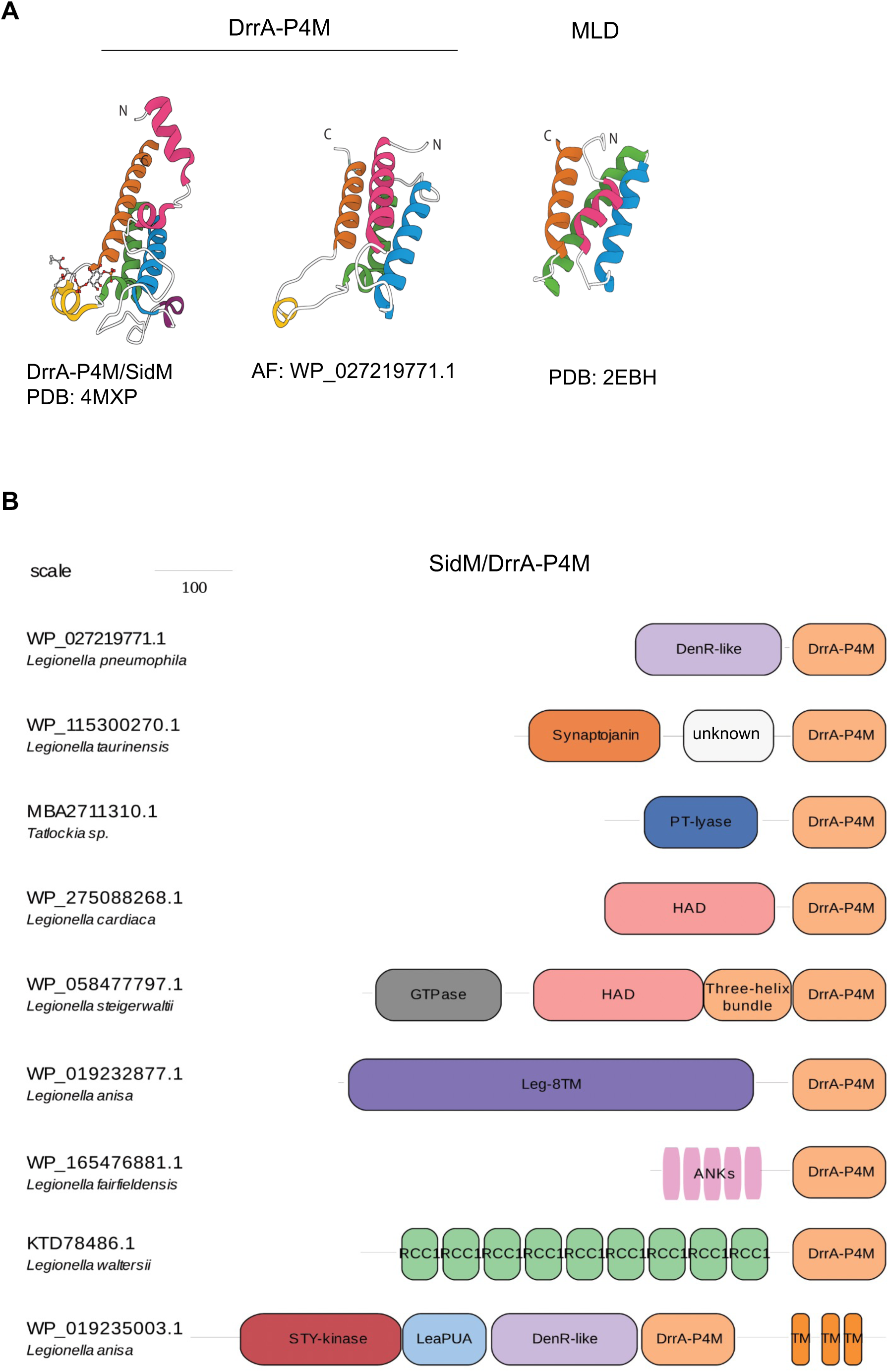
**(A)** Structures of SidM-P4M and MLD folds; experimentally determined structures are indicated by PDB identifiers, and predicted models correspond to Alphafold 3 entries. **(B)** Legionella effector architectures centered on the SidM-P4M domain.

**Supporting Fig. 5:**
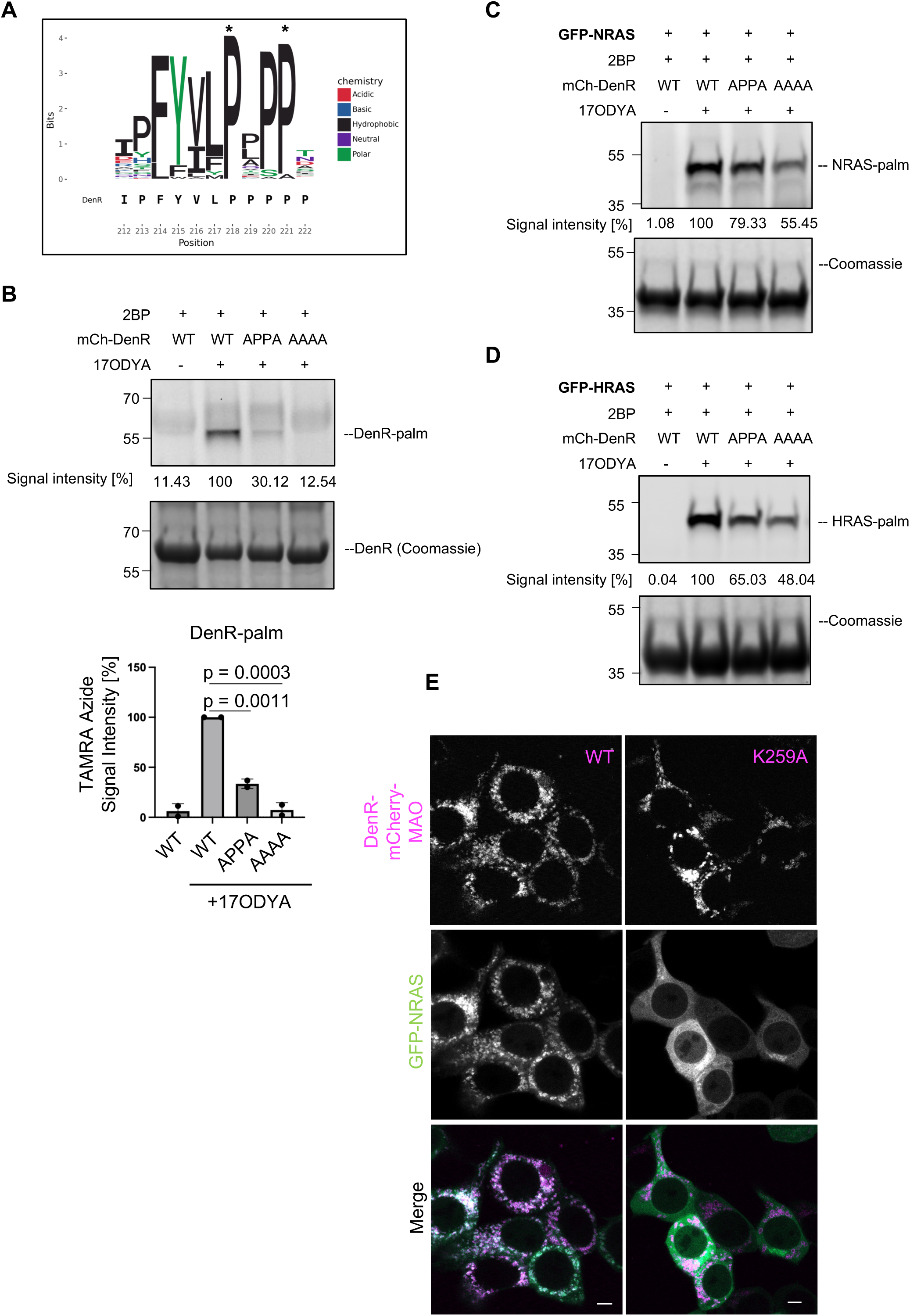
(**A**) Sequence Logo of conserved relay residues within the linker region of DenR. Lp02 DenR sequence is displayed along the bottom of the sequence logo, with amino acid positions numbered underneath. A patch of five prolines appears from aa218-222. Residue conservation across DenR homologs is displayed by character size in the sequence logo, with larger characters being more strictly conserved residues. The most highly conserved residues in this region, P218 and P221 (shown structurally in Figure 5F), are highlighted here with asterisks. (**B**) Click chemistry to detect the role of conserved linker residues on DenR autoacylation. 2BP-treated HEK293 cells producing the indicated mCherry-tagged DenR variants were metabolically labelled with 17ODYA. mCherry-tagged DenR was affinity-purified by immunoprecipitation with mCherry nanobody beads, and palmitoylation was detected by click chemistry with TAMRA-azide and in-gel visualization using a ChemiDoc Imaging System. Graphs represent mean +/− SD quantification of TAMRA-azide signal from two experiments (n=2). (**C,D**) Representative blots from Figure 5G and 5H quantifications, respectively. (**E**) NRAS recruitment by DenR requires PI3P binding. Same experiment as Figure 5I, but in HEK cells which display more consistent expression. All scale bars, 5 µm.

**Supporting Fig. 6:**
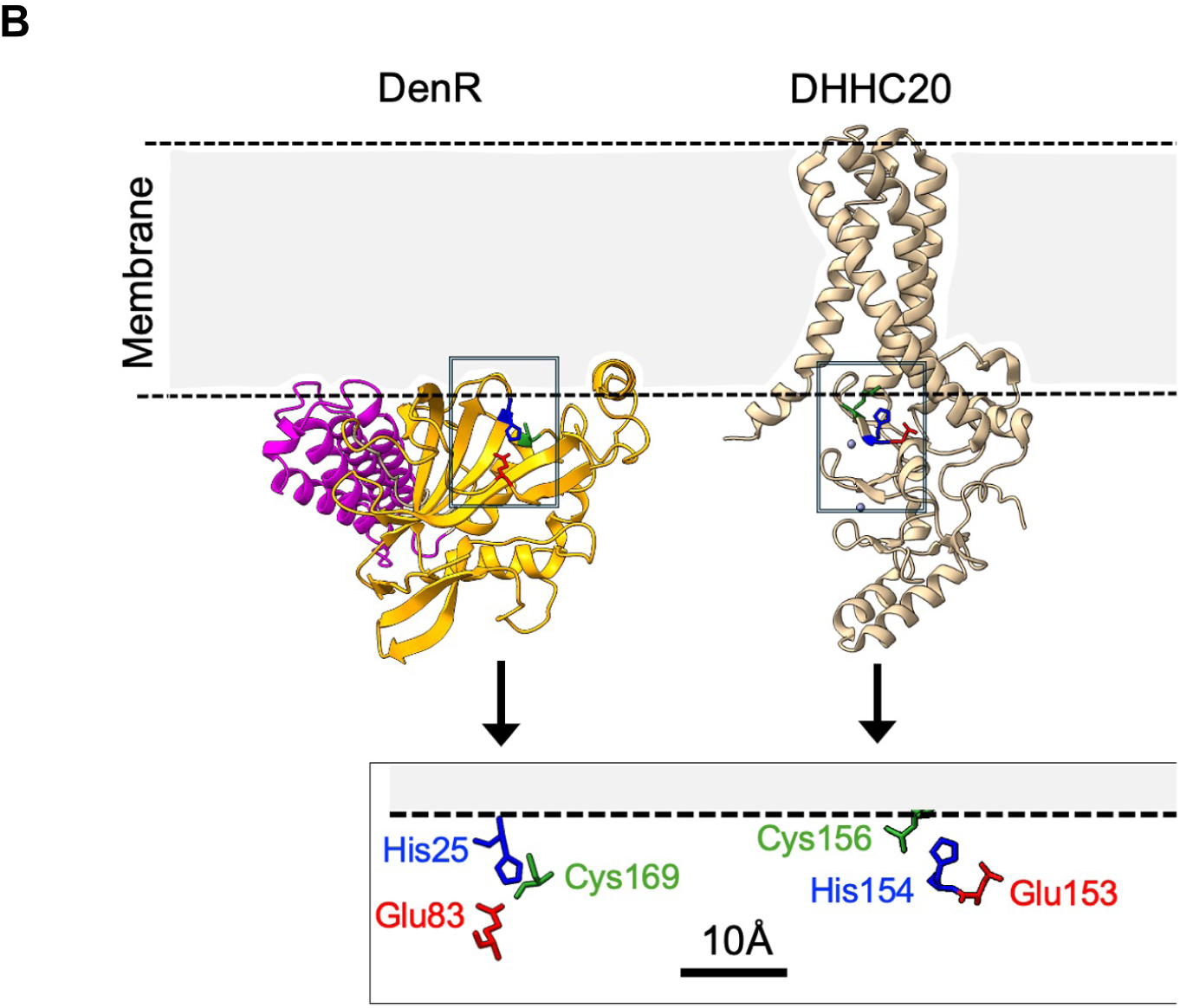
(**A**) Membrane binding model. DenR was placed onto a theoretical membrane, with the P4M domain and the cap domain acting as main anchor points. The DenR catalytic residues are shown in isolation relative to the lipid-water interface and are compared side-by-side to those from DHHC20 placed in the same membrane.

